# Inherent Specificity and Variation Sensitivity as Quantitative Metrics for RBP Binding

**DOI:** 10.1101/2025.03.28.646018

**Authors:** Soon Yi, Shashi S. Singh, Xuan Ye, Rohan Krishna, Vidusha Kothwela, Eckhard Jankowsky, Joseph M. Luna

## Abstract

RNA-binding proteins (RBPs) regulate every aspect of post-transcriptional gene expression, yet our ability to compare how selectively different RBPs recognize their targets remains limited. Binding affinity, expressed as a dissociation constant, provides a universal quantity for comparing binding strength, but no equivalent metric exists for binding specificity. Here we introduce two quantitative metrics to fill this gap: inherent specificity, which measures how selectively an RBP distinguishes its strongest binding motif from all other sequences, and variation sensitivity, which measures tolerance to single nucleotide changes within that motif. Analyzing high-throughput sequencing data across 100 RBPs *in vitro* and 27 in cells, we find strong correspondence between in vitro and cellular measurements for sequence-driven RBPs. Domain swap CLIP experiments demonstrate that specificity can be transferred between protein contexts. Mathematical modeling and cellular competition experiments reveal that low-specificity RBPs can paradoxically sharpen the target discrimination of high-specificity partners by occupying non-preferred sites, an emergent property not predictable from affinity alone. These metrics and accompanying R packages provide a practical framework for comparing RBP binding behaviors and modeling how RBPs compete for RNA targets across the transcriptome.

## Introduction

Gene expression in higher eukaryotic cells involves coordinated interactions between thousands of RNA-binding proteins (RBPs) and tens of thousands of RNAs^1–5^. These interactions create regulatory networks responsible for controlling all facets of post-transcriptional regulation of gene expression, including RNA splicing, export, translation, and decay^6–8^. Deregulation of RNA-RBP networks has been linked to a spectrum of human diseases, ranging from neuropathies and inflammations to various cancers^9–11^. Understanding the principles that govern RNA-RBP interactions is therefore central to understanding gene regulation in health and disease.

RNA-RBP interactions in eukaryotes are mediated by structurally diverse RNA-binding domains (RBDs), of which many functionally distinct classes have been discovered and characterized over the past few decades^1–3,12,13^. The RNA recognition motif (RRM) is the most prevalent, represented in approximately 2% of the human proteome^14^, while K-homology (KH) domains^15^, helicase domains^16^, dsRNA-binding domains^17^, PUF domains^18^, and many others further expand the structural repertoire available for RNA recognition^12,19^. Individual RBPs rarely rely on a single domain in isolation: multiple RBDs cooperate within a single protein or across protein complexes to achieve RNA binding^20^, and this modularity expands the RNA target space available to any given RBP. In addition to well-defined domain architectures, many RBPs harbor intrinsically disordered regions that engage RNA through multivalent, transient contacts^12,13,21^. Further, proteomic surveys have identified hundreds of additional proteins with no annotated RBD that nonetheless crosslink to RNA, including metabolic enzymes and cytoskeletal proteins whose RNA contacts remain poorly characterized^2,3,13^. How RBPs with such varied architectures and binding modes achieve selective RNA recognition remains incompletely understood.

Characterizing RNA-RBP interactions requires quantifying two distinct binding properties: affinity and specificity. Affinity, expressed as a dissociation constant (*K_d_*), measures the strength of an RBP-RNA interaction and provides a tractable quantity comparable across any two proteins for a given RNA substrate, as measured by biochemical assays such as EMSA, ITC, and SPR^22–25^. Decades of such measurements have established that RBPs bind their targets with affinities spanning picomolar to micromolar ranges^12^.

Affinity toward individual sequences, however, does not capture how an RBP binds to the full range of RNAs it encounters. High-throughput *in vitro* assays such as RNACompete^26,27^, RNA Bind-n-Seq^28,29^, and HITS-Eq^30,31^ have made it possible to measure binding across millions of sequences *in vitro*, and multiple CLIP-based approaches have extended this view to the cellular context, mapping transcriptome-wide binding patterns of hundreds of RBPs across diverse biological contexts^32–36^.

For *in vitro* binding data, the affinity landscape is typically shown as a preferred motif, while CLIP data are interpreted in terms of target occupancy. In both cases, binding behavior is classified as a binary distinction of specific versus non-specific. Tools such as sequence logos and position weight matrices formalize this, capturing what an RBP prefers to bind but not how strongly it discriminates that preference from all other potential binding sites^37,38^. Unlike affinity, for which Kd provides a universal and comparable quantity, no equivalent metric exists for binding specificity, limiting systematic comparison across RBPs and impeding mechanistic modeling of how they compete for targets in cells. Related distributional analyses of RBNS enrichment profiles have been explored in the context of individual RBPs^39^, but have not been condensed into standardized cross-RBP metrics suitable for systematic ranking or predictive modeling.

Here, we address this gap by introducing two quantitative metrics for RBP binding specificity. Inherent specificity (IS) measures how selectively an RBP binds its top motif relative to all other sequences of the same length. Variation sensitivity (VS) measures tolerance to single nucleotide changes within that top motif. Analyzing RBNS datasets across 26 single-domain RBPs, we show that IS and VS are robust across motif lengths and inter-correlated. Comparing these *in vitro* metrics to cellular binding data from eCLIP, we find that correspondence between inherent and cellular specificity depends on the local RNA structural context of target motifs, with sequence-driven RBPs showing strong agreement across both contexts. To test the extent to which specificity is a portable property of an RRM, we performed domain-swap CLIP experiments between high-specificity HNRNPC and low-specificity RBM25, demonstrating that RRM identity contributes to, but does not fully determine, cellular binding specificity. We further show that IS and VS predict how low-specificity RBPs can enhance the target discrimination of high-specificity RBPs by occupying non-target sites. We experimentally confirm this phenomenon with *in vitro* iCLIP data and validate in cells through HNRNPC CLIP experiments in the presence of non-specific RBM25. Together, these findings establish IS and VS as quantitative descriptors of RBP binding specificity, providing a basis for comparing RBPs and modeling RNA target competition in complex cellular environments.

## Results

### Inherent specificity and variation sensitivity as distinct metrics to quantify RNA-RBP interactions

To understand how RBPs interact with their RNA targets, we consider the full distribution of binding affinities across all possible sequence variants (motifs) of a given K-mer (**Fig. 1A, left**). Affinity and specificity are distinct properties of this distribution. While affinity describes the absolute strength of binding to a given sequence, typically expressed as a dissociation constant (*K_d_*), specificity describes the selectivity of an RBP for its preferred sequence, compared to other sequences. A highly specific RBP binds to a narrow set of sequence variants while showing significantly reduced binding to all other sequence variants. An RBP with little specificity, on the other hand, binds many sequence variants similarly. (**Fig. 1A, middle**). Of note, an RBP with high affinity for its preferred sequence may still bind many other sequences nearly as well, while an RBP with modest absolute affinity may discriminate its preferred motif from all others (**Fig. 1A, right**). This distinction between affinity and specificity is well established for protein-DNA interactions^38^ and applies to protein-RNA interactions^37^. Yet while affinity for a given sequence can be directly compared across any two proteins via *K_d_*, no equivalent quantitative descriptor exists for specificity.

**Figure 1:**
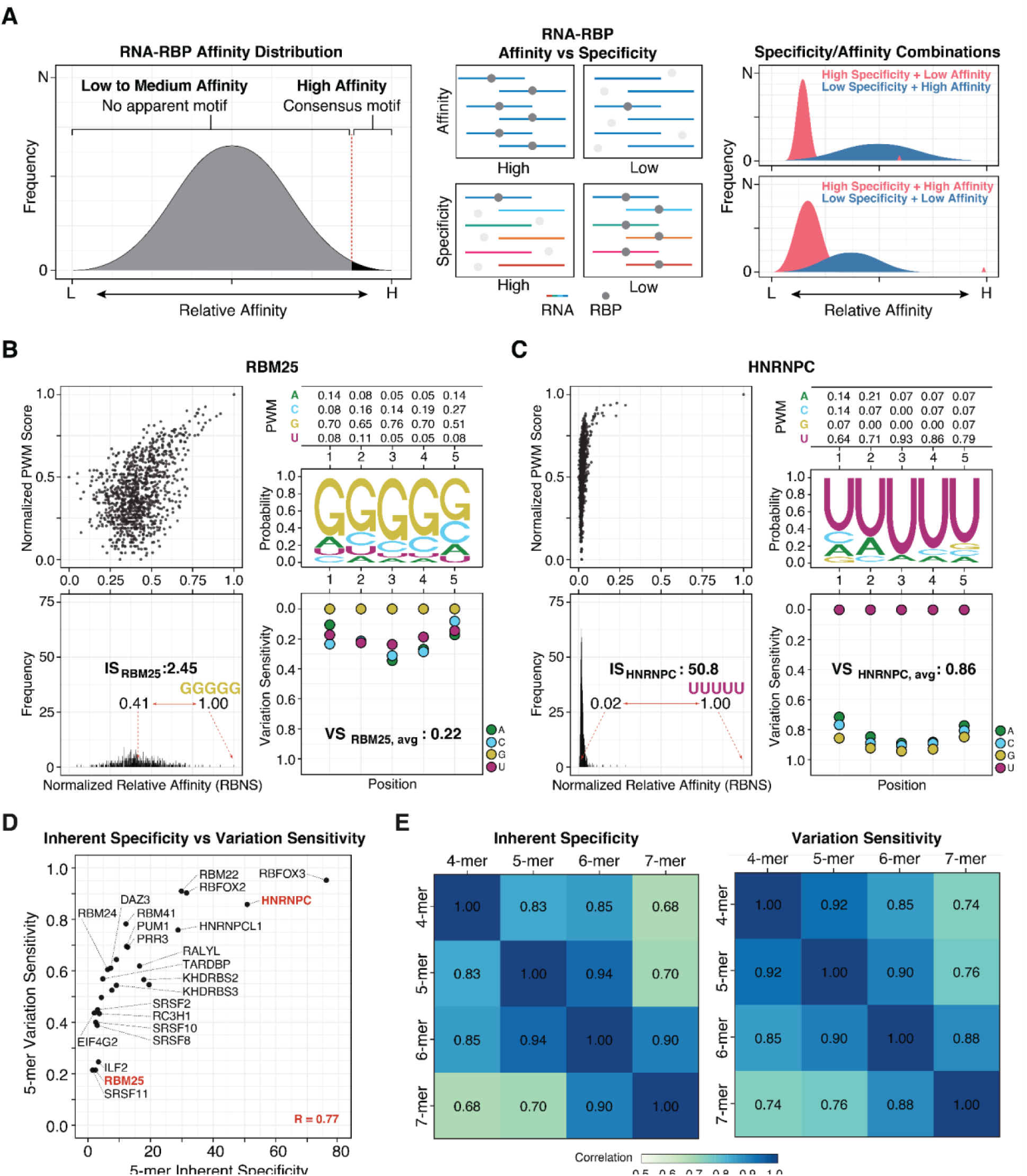
Inherent specificity (IS) and variation sensitivity (VS) to quantify RNA binding protein behavior. **(A)** The relative affinity distribution depicts the RNA binding landscape of an RBP. High-affinity motifs are used for consensus motif generation (left). Binding affinity describes the strength of the binding, while specificity describes the selectivity of an RBP towards its target motif (middle). These metrics do not always correlate, resulting in four possible categories of RBP binding profiles (right). **(B)** Graphical characterizations of the 5-mer RNA binding behavior of RBM25. A comparison of measured relative affinity distributions (from RBNS; x-axis) and reconstituted relative affinity distributions (from the PWM; y-axis) is shown (top left). The position weight matrix (PWM), derived from RBNS 5-mer relative affinity distribution, and the sequence logo, a graphical representation of the PWM, illustrate the traditional approach to characterizing the binding behavior of RBM25. Inherent specificity (bottom left) measures the affinity ratio between the top and the median binding motifs. Variation sensitivity (bottom right) describes the average deviation in binding affinities to 5-mer sequence motifs with a single nucleotide change from the top binding motif (GGGGG). **(C)** The same as **(B)** but for HNRNPC. **(D)** Correlation between 5-mer IS and VS for 26 single-domain RBPs from RBNS data. The two representative RBPs, RBM25 (low specificity) and HNRNPC (high specificity), are highlighted in red. Pearson correlation is shown at the bottom right in red. **(E)** Pairwise Pearson correlations across different *K*-mers for IS (left) and VS (right) ranks for the 26 RBPs shown in **(D)**.

To illustrate this, we analyzed RBNS binding data from the ENCODE consortium^29,40^ for RBM25 and HNRNPC, two RBPs each containing a single RRM but with distinct binding preferences (**Fig. 1B-C**). Sequence logos derived from their top 5-mer motifs confirm that both proteins are moderately motif-specific, with RBM25 binding G-rich motifs and HNRNPC preferring U-rich motifs (**Fig. 1B-C, top right**). PWMs offer a more quantitative representation of the same information and can be used to reconstruct predicted motif enrichment distributions^41^, which we compare directly to measured RBNS R-scores as a proxy for relative binding preference^42^ (**Fig. 1B-C, top left;** see Methods). While RBNS R-scores do not directly measure *K_d_* values, they have been shown to correlate strongly with biophysically measured *K_d_* values^28^, supporting their use as a proxy for relative binding affinity. To ensure comparability across RBPs, we normalize R-scores to a common scale prior to analysis (see Methods). This comparison reveals that PWM-predicted scores can diverge substantially from RBNS, where here the discrepancy is most pronounced for HNRNPC. The source of this divergence becomes clear when examining the full relative affinity distributions: HNRNPC’s distribution is bimodal, with strong preference for UUUUU and lower relative affinity for all other sequences, while the RBM25 distribution is broad and bell-shaped (**Fig. 1B-C, bottom left**). While sequence logos and PWMs suggest comparable specificity for the two proteins, their measured RBNS relative affinity distributions suggest otherwise.

To address this gap, we introduce two metrics that quantify RBP binding specificity from relative affinity distributions to enable direct comparison across RBPs. The first, *inherent specificity* (IS), measures how selectively an RBP binds its top motif relative to all other sequences of the same length. IS is defined as the ratio of the affinity for the top binding motif to the affinity of the median motif, where the median serves as a representative baseline for the full distribution. Applied to RBM25 and HNRNPC, IS reveals that HNRNPC is approximately 20-fold more specific than RBM25, a difference invisible to PWMs and sequence logos (**Fig. 1B-C, bottom left**).

The second metric, *variation sensitivity* (VS), captures a complementary property: how sensitively an RBP discriminates between its top motif and all single-nucleotide variants of that motif (**Fig. 1B-C, bottom right**). VS is calculated as the normalized relative affinity of each single-nucleotide variant relative to the top binding motif, averaged across all variant positions. Like IS, VS enables direct cross-RBP comparison, showing for example that HNRNPC is approximately four times more sensitive than RBM25 to substitutions in its preferred motif. A high VS value indicates that single-nucleotide deviations from the top motif produce substantial drops in enrichment, distinguishing true motif preference from enrichment driven by subsequence overlap with the top motif.

### Inherent specificity and variation sensitivity show strong inter- and intra-correlations across different *K*-mers

To understand how changes in *K*-mer length affect these metrics, we calculated IS and VS values from 4 to 7-mers for RBNS datasets (**Table S3**), focusing on the 26 RBPs with a single RNA binding domain purified for the RBNS assay^29^ to exclude confounding effects from multi-RBD interactions. Comparing IS and VS values across *K*-mer lengths, we observed that higher IS RBPs tended to be sensitive to changes in their top motifs (**Fig. 1D, Fig. S1A**). For all 26 RBPs, IS values increased with K-mer length (**Fig. S1B, left**), with the most pronounced increases in higher-specificity RBPs: PRR3 (6-mer IS: 30.8, 7-mer IS: 96.6), RBFOX3 (4-mer IS: 17.2, 5-mer IS: 76.1), and RBM22 (5-mer IS: 29.9, 6-mer IS: 93.9) each showed significant jumps at specific *K*-mer length transitions. VS values were more stable across K-mer lengths, with PRR3, KHDRBS3, DAZ3, RBM24, and EIF4G2 as the primary exceptions (**Fig. S1B, right**). Importantly, despite these absolute changes, the rank order of both IS and VS across RBPs was preserved across all K-mer lengths tested (**Fig. S1C, Fig. 1E**). That rank order is preserved despite absolute IS values increasing with k-mer length indicates that these metrics reflect stable properties of each RBD rather than artifacts of the sequence length used to measure them. Based on this, we used 5-mer calculations for subsequent analyses.

### Inherent and cellular binding metrics diverge in a structure-context-dependent manner

We next compared IS and VS values derived from *in vitro* RBNS data with RBP binding activities in cells. RBNS measures inherent binding properties by exposing a single RBP to an equal mix of all possible K-mer substrates^28^, a controlled condition that differs substantially from the cellular environment, where competing RBPs, variable expression levels, tissue-specific factors, and subcellular compartmentalization all shape RNA target availability.

To compare inherent binding metrics to cellular binding behavior, we calculated motif enrichment scores for all 5-mers from eCLIP datasets for 28 RBPs with corresponding RBNS data (**Table S1**, Methods), terming these ‘cellular’ specificity (CS) and variation sensitivity (CVS) to distinguish them from ‘inherent’ metrics derived from RBNS. To illustrate the range of behaviors observed, we highlight four RBPs spanning a broad range of IS values: EIF4G2, PCBP2, RBFOX2, and HNRNPC (**Fig. 2A**); EIF4G2 was used in place of RBM25, for which no eCLIP data are available.

**Figure 2:**
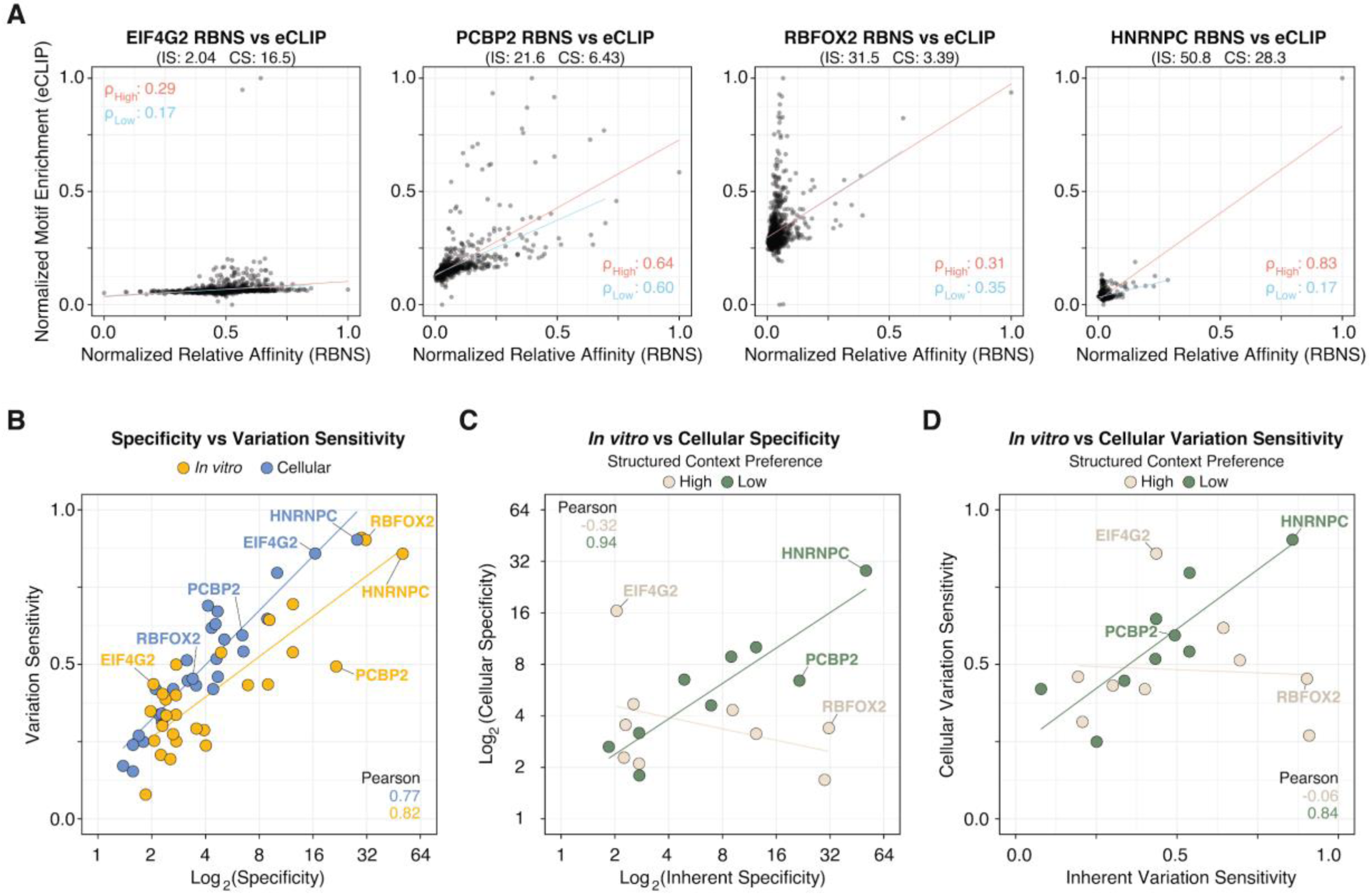
Comparison of inherent and cellular specificity and variation sensitivity shows a structure-context-dependent correlation. **(A)** Correlation between 5-mer RNA-RBP motif enrichments measured by RBNS (*in vitro*) and eCLIP (cellular). ρ_High_ and ρ_Low_ indicate Pearson correlation coefficients for the full motif enrichment (1,024 5-mer motifs) and the low-enrichment motif subsets without the top 10 5-mer motifs (1,014 5-mer motifs), respectively. Inherent specificity (IS) from RBNS and cellular specificity (CS) from eCLIP are shown for each RBP. **(B)** Relationship between specificity (log_2_) and variation sensitivity for *in vitro* (RBNS, yellow) and cellular (eCLIP, blue) datasets. Selected RBPs from **(A)** are labeled. **(C)** Comparison of Inherent Specificity (log_2_) versus Cellular Specificity (log_2_). Points are colored by structured context preference (High vs. Low). Pearson correlation coefficients for each group are indicated at the bottom right. **(D)** Correlation between Inherent Variation Sensitivity and Cellular Variation Sensitivity, categorized by structured context preference as in **(C)**.

EIF4G2 (IS: 2.04) showed poor agreement between IS and CS, while PCBP2 (IS: 21.6), roughly 10-fold more specific *in vitro*, showed higher correlation. For RBFOX2 (IS: 31.5) and HNRNPC (IS: 50.8), the top motifs identified *in vitro*, GCAUG and UUUUU respectively, remained dominant in cells (**Fig. 2A**). However, RBFOX2 specificity decreased more than 9-fold in cells (IS: 31.5, CS: 3.39), while EIF4G2 increased nearly 8-fold (IS: 2.04, CS: 16.5), demonstrating that *in vitro* IS values are not sufficient to predict the degree or direction of CS shifts (**Fig. 2A**). Specificity and variation sensitivity were positively correlated within each context, but the rank order of RBPs differed between *in vitro* and cellular measurements (**Fig. 2B**).

To understand what may drive these discrepancies, we grouped RBPs based on their known preferences for RNA secondary structure context around the target motif^29^. RBPs with increased enrichment in stem loops, stem bulges, or stems were designated as having high structural context preference; all others were designated as having low structural context preference. We observed little correlation between IS and CS for RBPs that recognize RNA motifs in a local structural context (Pearson r = −0.32; **Fig. 2C**). In contrast, RBPs with a low structural context preference showed a strong positive correlation between IS and CS (Pearson r = 0.94), and a similarly strong correlation between VS and CVS (Pearson r = 0.84; **Fig. 2D**). Together, these results indicate that structural context is a key determinant of whether inherent binding properties are preserved in cells: when binding depends on local RNA structure, *in vitro* and cellular metrics diverge substantially, while sequence-driven RBPs show strong correspondence across both experimental contexts.

### Inherent specificity encoded in RRMs is transferable between protein contexts

Our comparison of *in vitro* and cellular measurements revealed that for RBPs with low structural context preferences, such as HNRNPC, inherent and cellular binding metrics correspond closely. However, RBNS experiments characterize isolated RNA binding domains while CLIP captures full-length proteins in their native cellular environment. To determine how much of an RBP’s cellular binding specificity is encoded within its RRM, we designed an RRM-swap experiment between two single-RRM RBPs with contrasting specificities: HNRNPC (IS: 50.8), which binds U-rich motifs with high specificity, and RBM25 (IS: 2.04), which binds G-rich motifs with low specificity. Despite structural similarity, the two RRMs share only 18% sequence identity (**Fig. S1D-E**).

We engineered HEK293 cells to stably express V5-tagged wild-type and RRM-swapped versions of HNRNPC and RBM25 under doxycycline induction (**Fig. 3A, Fig. S2A-B**). While numerous CLIP and eCLIP datasets exist for HNRNPC^34,43–45^, no CLIP data exist for RBM25. RNA-IP experiments have suggested it binds G-quadruplex structures^46^, consistent with the low sequence specificity seen in RBNS data (**Fig. 1B**). After confirming that V5-tagged proteins localized to the nucleus as expected for both endogenous proteins (**Fig. S2C**), we performed CLIP on all four cell lines (**Fig. 3A, Fig. S2D**) and analyzed libraries using the CLIPittyClip pipeline^47^, with reanalyzed wild-type HNRNPC CLIP and eCLIP datasets^40,43^ processed in parallel as analysis controls.

**Figure 3:**
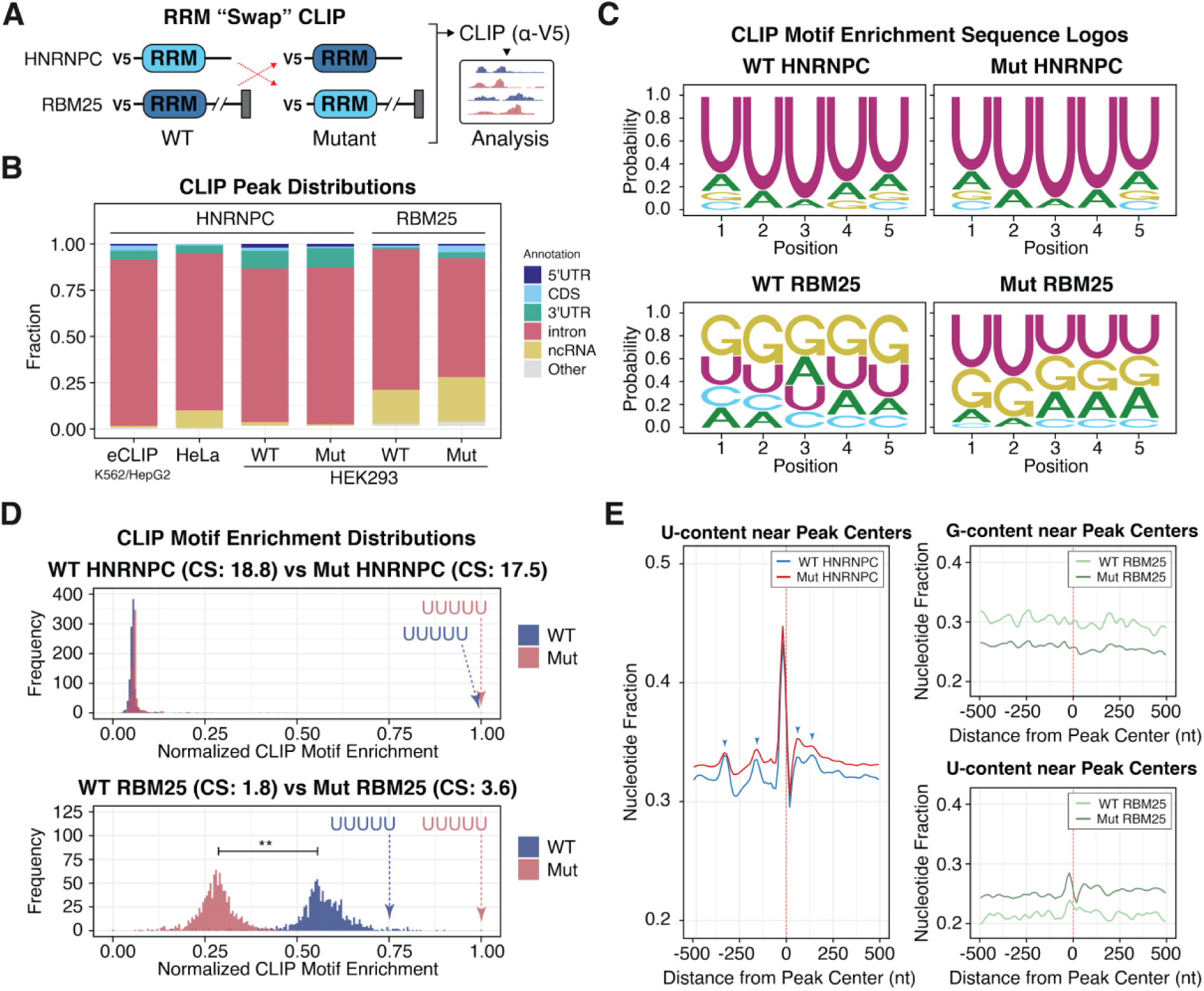
RRM swap between HNRNPC and RBM25 reveals that Inherent Specificity transfer is dependent on the protein-specific context. **(A)** Schematic of the RRM swap CLIP between HNRNPC and RBM25. **(B)** CLIP peak distributions across genomic regions for the indicated datasets. eCLIP data from K562 and HepG2 were merged for analysis. **(C)** Sequence logos derived from motif enrichment within CLIP peaks for wild-type (WT) and mutant (Mut) RBPs. **(D)** 5-mer motif enrichment distributions derived from CLIP peaks. Cellular specificity (CS) values are indicated for WT and Mut pairs, showing a significant increase for RBM25 (1.8 to 3.6) but stability for HNRNPC (18.8 to 17.5). The relative position of the “UUUUU” motif is indicated by dashed arrows. Statistical significance between distributions was determined by a two-tailed t-test (**, p < 0.01). **(E)** Nucleotide frequency analysis near CLIP peak centers. For HNRNPC (left), U-content for WT (blue) and Mut (red) is shown; periodic U-enrichment peaks are indicated by blue markers. For RBM25 (right), G-content (top) and U-content (bottom) near peak centers are shown for WT (light green) and Mut (dark green).

CLIP peak annotations were consistent across HNRNPC datasets, with strong intronic enrichment in both wild-type and mutant (**Fig. 3B**). RBM25 maintained its intronic and ncRNA preferences in both wild-type and mutant contexts. The RRM swap had minimal effect on genomic region preferences for either protein.

Sequence motif preferences were affected differently between the two proteins (**Fig. 3C**). Mutant HNRNPC maintained U-rich motif binding comparable to wild-type, with little change despite the RRM swap. Mutant RBM25, by contrast, showed decreased preference for G-rich motifs and increased preference for U-rich motifs, consistent with the introduction of the HNRNPC RRM. While PWMs suggested comparable motif specificity for both proteins, motif enrichment distributions revealed stark differences (**Fig. 3D**). Wild-type and mutant HNRNPC showed nearly identical cellular specificity for UUUUU (CS: 18.8 and 17.5, respectively), while mutant RBM25 showed a significant 2-fold increase relative to wild-type (CS: 1.8 and 3.6, respectively). The mutant RBM25’s specificity for UUUUU increased while its specificity for other U-rich motifs generally decreased (mean fold-change: 0.59, **Table S4**), recapitulating HNRNPC’s characteristic preference for UUUUU over other poly-U motifs.

HNRNPC is well known to form homotetramers^48–51^, raising the possibility that mutant subunits form heterotetramers with endogenous wild-type protein, potentially obscuring the effect of the RRM swap. Previous iCLIP results with HNRNPC described stereotypical periodicity of U-content approximately every 100-200 nucleotides near binding sites, attributed to oligomerization-driven spacing along the RNA^34^. We observed the same periodicity in our wild-type HNRNPC CLIP data, but found it visibly relaxed in the RRM-swapped mutant (**Fig. 3E**), indicating that the swap does alter HNRNPC engagement with poly-U stretches even in the presence of endogenous protein. We further note that the same analysis with wild-type and mutant RBM25 revealed a reciprocal pattern: wild-type RBM25 showed G-enrichment near peak centers with no apparent U periodicity, while the mutant showed reduced G-enrichment and an emerging U-content peak (**Fig. 3E**), consistent with partial acquisition of HNRNPC-like sequence recognition. To minimize tetramerization with endogenous protein directly, we knocked down endogenous HNRNPC by greater than 80% in our stable cell lines (**Fig. S2E**) and repeated V5 IP-CLIP for both wild-type and mutant constructs (**Fig. S2F**). Peak annotations, sequence logos, and motif enrichment distributions were unchanged relative to the non-knockdown condition, and the characteristic U-content periodicity observed in wild-type HNRNPC was preserved and again visibly relaxed in the mutant (**Fig. S3A-D**). While endogenous protein was not completely eliminated, these data together suggest that some poly-U specificity is encoded within the HNRNPC RRM, but the full extent of HNRNPC’s sequence selectivity in cells involves additional determinants beyond the RRM alone.

Cellular VS values showed corresponding shifts. HNRNPC maintained similar values across wild-type and mutant constructs (0.80 and 0.78, respectively), while mutant RBM25 showed a notable increase relative to wild-type (0.41 to 0.61), reflecting increased sensitivity to substitutions in the UUUUU motif consistent with partial adoption of HNRNPC-like binding behavior (**Fig. S3E**). These effects are further evident in sequence context logos showing nucleotide frequencies near peak centers, where the mutant RBM25 displays a broader U-rich context that resembles wild-type HNRNPC, while mutant HNRNPC is unchanged (**Fig. S3G-J**). Taken together, the directional shift in RBM25 and the consistency of HNRNPC binding confirm that CS and CVS are responsive to engineered changes in RBP sequence recognition.

### Inherent specificity and variation sensitivity shape competitive RNA binding

Our RRM-swap experiments showed that binding specificities can be partially transferred between proteins by exchanging their RRMs, raising the question of how different combinations of IS and VS values influence competitive RNA binding when multiple RBPs target the same sequences. To explore this, we developed a mathematical model of four hypothetical RBPs with varying IS and VS values interacting with a single target RNA (**Fig. 4A**). All four RBPs were designed to bind uridine-rich motifs with the highest affinity and guanine-rich motifs with the lowest, but with distinct binding landscapes defined by their IS (**Fig. 4B**) and VS values (**Fig. 4C**). For the simulation, all four RBPs were introduced simultaneously to ensure equal access to the RNA, and each binding site was treated as an independent event. This allowed us to calculate equilibrium concentrations of RNA-RBP complexes at each binding site (**Fig. 4D**), which were then used to derive the binding probability of each RBP at a given target site (see Methods).

**Figure 4:**
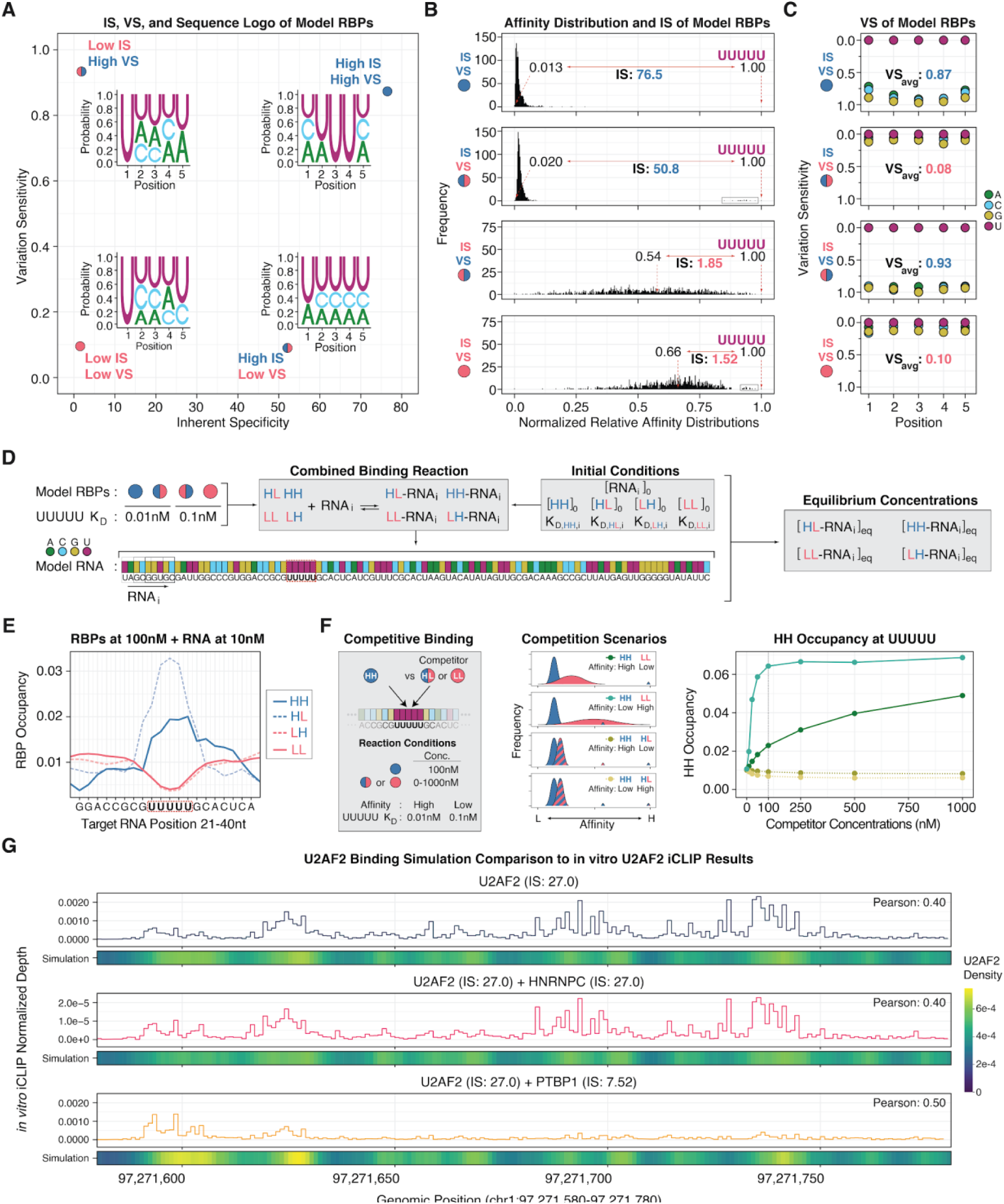
*In silico* modeling of RBP binding using model RBPs and *in vitro* binding data shows that non-specific RBPs focus the binding of high-specificity RBPs. **(A)** Classification of four model RBPs by their Inherent Specificity (IS) and Variation Sensitivity (VS) values: HH (High IS, High VS), HL (High IS, Low VS), LH (Low IS, High VS), and LL (Low IS, Low VS). Sequence logos for each model are shown. **(B)** Affinity distributions for the four model RBPs, highlighting median and top affinity values and the resulting IS. **(C)** VS plots and average VS values for the four model RBPs. **(D)** Schematic of the mathematical binding model. Dissociation constants (*K_d_*) for the target motif “UUUUU” are indicated; *K_d_* values for all other 5-mers are derived from the respective affinity distributions. Equilibrium concentrations are used to calculate per-position binding occupancies on the model RNA. The target binding site is marked in red boxes. **(E)** Simulated occupancies for model RBPs at equimolar concentrations (100 nM) across the model RNA (10 nM). **(F)** Competition simulation of HH binding at the target site in the presence of either LL or HL competitors. Four scenarios vary the affinity and specificity of the competitor. HH occupancy at the target site across a range of competitor concentrations is shown (right). **(G)** Simulation of U2AF2 binding to the *PTBP2* intronic region compared to *in vitro* iCLIP results. Simulations show U2AF2 alone (top), with HNRNPC (high specificity, middle) or with PTBP1 (low specificity, bottom). Line plots represent iCLIP enrichment signals; heatmaps depict simulated U2AF2 occupancy. Modeling utilized 7-mer data from RNACompete.

When all four model RBPs were simulated at equimolar concentrations, high-specificity RBPs (HH and HL, modeled with 0.01nM UUUUU *K_d_*) showed preferential occupancy at the UUUUU target site, while low-specificity RBPs (LH and LL, modeled with 0.1nM UUUUU *K_d_*) distributed across the RNA, accumulating at non-preferred sites and avoiding UUUUU (**Fig. 4E, Fig. S4A**). We interpret this to mean that by occupying non-target sites, low-specificity RBPs reduce the search space available to high-specificity RBPs. In this scenario, any low-specificity RBP that does occupy a preferred site of high specificity RBP will be competitively displaced.

To model this directly, we simulated HH occupancy at UUUUU in the presence of either a low-specificity (LL) or high-specificity (HL) competitor across a range of concentrations and, importantly, simulated binding affinities (**Fig. 4F, Fig. S4B-C**). When LL had lower affinity for UUUUU than HH, increasing LL concentration enhanced HH occupancy at the target, consistent with LL sequestering non-target sites. When LL affinity exceeded that of HH, enhancement of HH occupancy at UUUUU was still observed, as high affinity for all sites caused LL to distribute across the RNA, further reducing competition at UUUUU. In contrast, HL competitors suppressed HH occupancy at UUUUU regardless of relative affinity, consistent with direct competition for the same target site. Together, these simulations show that whether a competing RBP enhances or suppresses target site occupancy depends on its specificity profile, not on affinity alone.

To test whether these predictions extend to real RBPs, we sought an experimental dataset in which a single protein was profiled by CLIP in the presence or absence of a competitor. Sutandy et al. performed *in vitro* iCLIP on U2AF2 in the presence or absence of competing RBPs HNRNPC and PTBP1^52^, providing a direct test of our predictions. U2AF2, PTBP1, and HNRNPC all bind poly-pyrimidine-rich motifs, and using RNACompete datasets^27^ we calculated that U2AF2 (IS: 27.0) and HNRNPC (IS: 27.0) are each approximately four times more specific than PTBP1 (IS: 7.5). We therefore predicted that low-specificity PTBP1 would enhance U2AF2 binding to its preferred motif sites, while equally specific HNRNPC would not, and simulated U2AF2 binding to the intronic region of the PTBP2 transcript to compare against the iCLIP data. Simulating U2AF2 alone recapitulated its measured binding distribution across the PTBP2 intronic region (**Fig. 4G, top**). Adding HNRNPC as a competitor had little effect, with U2AF2 maintaining its distributed binding pattern, (**Fig. 4G, middle**). In contrast, adding PTBP1 sustained and focused U2AF2 binding at its principal target sites (**Fig. 4G, bottom**), increasing Pearson correlation between the measured and the predicted binding from 0.4 to 0.5. These results demonstrate that IS values derived from *in vitro* binding data can be used to model competitive binding outcomes under defined biochemical conditions.

### A low-specificity competitor focuses HNRNPC binding toward preferred sites in cells

The *in vitro* iCLIP results encouraged us to ask whether the same competitive logic could be observed in cells. Having already established HNRNPC and RBM25 CLIP in our stable cell lines, we reasoned that performing HNRNPC CLIP in the presence of exogenously overexpressed RBM25 would provide a direct cellular test of our competition model, an approach we term “competition CLIP” (**Fig. 5A**). Based on our model, we predicted that overexpressing low-specificity WT RBM25 would focus HNRNPC binding toward its preferred U-rich sites by occupying non-target sites, while overexpressing mutant RBM25, with its increased U-rich specificity, would compete directly with HNRNPC.

**Figure 5:**
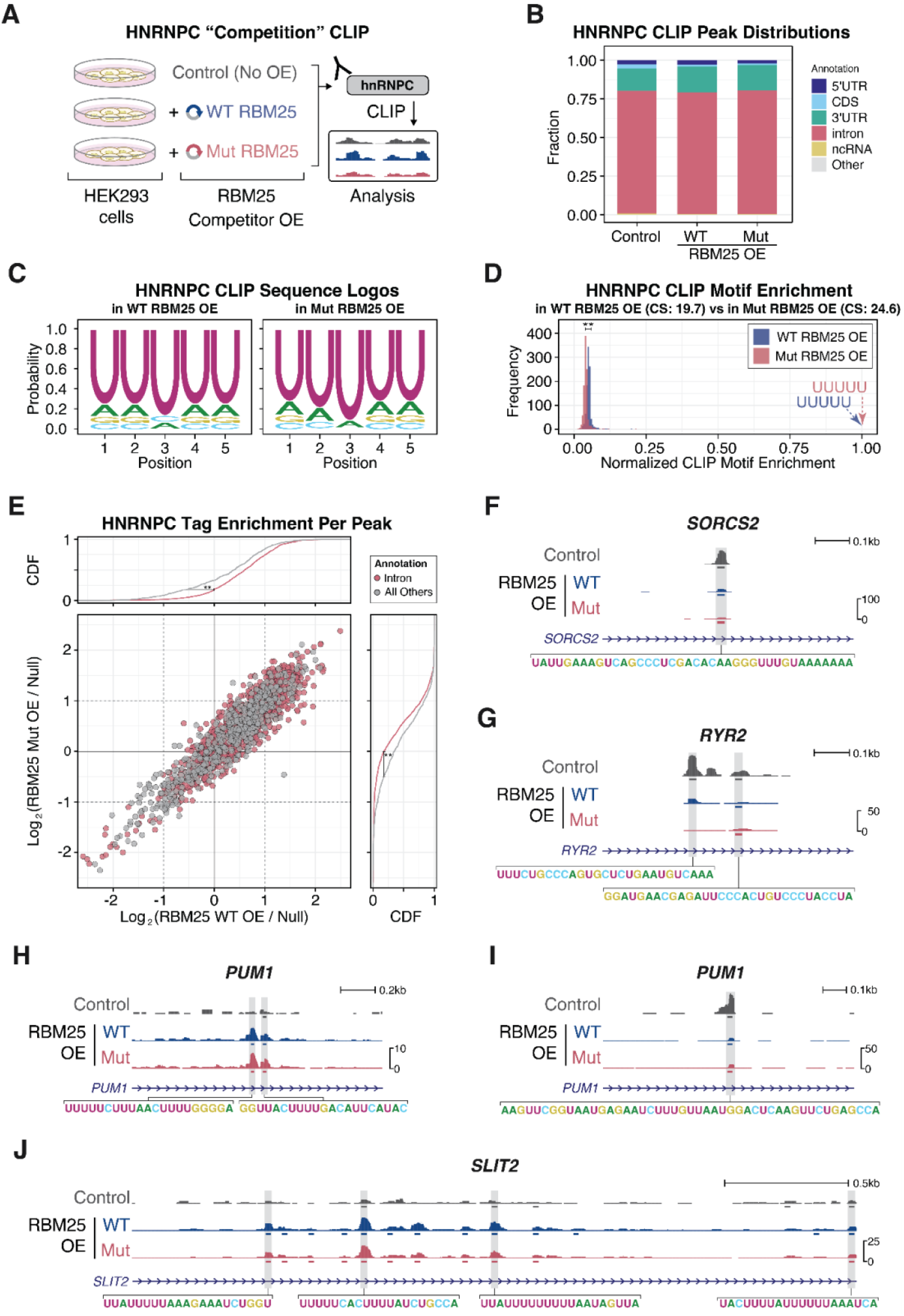
Cellular HNRNPC binding is shaped by competition with a non-specific RBP. **(A)** Schematic of HNRNPC competition CLIP during wild-type (WT) or mutant (Mut) RBM25 overexpression. **(B)** Genomic distribution of HNRNPC CLIP peaks across conditions. **(C)** HNRNPC sequence logos derived from CLIP motif enrichment. **(D)** 5-mer motif enrichment distributions for HNRNPC. Cellular Specificity (CS) values are shown for HNRNPC during WT RBM25 OE (19.7) and Mut RBM25 OE (24.6). Asterisks indicate statistical significance by a two-tailed t-test (**, p < 0.01). (**E**) Per-peak HNRNPC tag enrichment comparison between WT and Mut RBM25 overexpression conditions. Tag enrichments are calculated as the ratio of tag counts upon RBM25 overexpression to tag counts in the null, non-overexpressed condition. Points are colored by genomic annotation (Intron vs. All Others), with cumulative distribution functions shown on the marginal axes. Asterisks indicate statistical significance by a Kolmogorov-Smirnov test (**, p < 0.01) Depth-normalized genomic tracks for *SORCS2* **(F)**, *RYR2* **(G)**, *PUM1* **(H,I)**, and *SLIT2* **(J)** introns showing HNRNPC peak suppression and focusing effects upon RBM25 overexpression. Identified peaks are shown under each track. Nucleotide sequences within the peak regions are shown.

We performed HNRNPC V5 CLIP in cells overexpressing either WT or mutant RBM25 and compared peak distributions, motif preferences, and binding profiles to a no-overexpression control (**Fig. 5A-D**). Genomic region preferences (**Fig. 5B**) and U-rich motif preference (**Fig. 5C**) were unchanged across conditions. To anticipate the experimental outcome, we first simulated HNRNPC competition with either a UUUUU- or GGGGG-preferring low-specificity competitor using model RBPs, analogous to mutant and WT RBM25 (**Fig. S5A**), then repeated the simulation using actual HNRNPC and RBM25 CLIP motif enrichment data (**Fig. S5B**). Both predicted that WT RBM25 would focus HNRNPC toward U-rich sites, while mutant RBM25, with its modest gain in U-rich specificity, would be a less effective competitor at non-preferred sites. Consistent with this, HNRNPC cellular specificity increased from 19.7 to 24.6 under mutant RBM25 overexpression compared to WT (**Fig. 5D**).

Systematic comparison of per-peak HNRNPC tag enrichment across conditions revealed that the effects of WT and Mut RBM25 overexpression were similar in direction where peaks that gained HNRNPC binding under WT overexpression also gained binding under Mut overexpression, and vice versa (**Fig. 5E**). The most consistent changes were observed at intronic peaks, which showed a significant shift toward increased HNRNPC enrichment under both overexpression conditions (**Fig. 5E, Fig. S5C**). That both competitors produced similar focusing effects is consistent with both remaining low-specificity relative to HNRNPC, even accounting for the measurable increase in Mut RBM25 specificity (**Fig. 3D**).

Examination of individual loci illustrates the range of effects observed. At intronic peaks in SORCS2 (**Fig. 5F**), RYR2 (**Fig. 5G**), and PUM1 (**Fig. 5H**) that lacked poly-U stretches, we observed suppressed HNRNPC binding upon RBM25 overexpression, suggesting that non-preferred sites are masked by the presence of a low-specificity competitor. However, these effects were not unidirectional within a transcript as a separate PUM1 intronic region with poly-U stretches present within or near the peak region showed increased HNRNPC binding upon RBM25 overexpression (**Fig. 5I**), consistent with focusing. The SLIT2 locus illustrates that focusing can occur across multiple U-rich sites within a single transcript, with WT or mutant RBM25 overexpression enhancing HNRNPC binding at several intronic positions (**Fig. 5J**). As a whole, these results show that the competitive binding dynamics predicted by IS and VS values are observable in cells, and that the specificity profile of a competing RBP can shape the binding landscape of a higher-specificity partner through a combination of focusing or competitive masking of occupancy across target sites.

### IS and VS reveal specificity patterns across the cellular RBP repertoire

To examine how IS and VS scale across a broader set of RBPs in cells, we calculated CS and CVS from 251 eCLIP datasets covering 168 RBPs across two cell lines, HepG2 and K562 (**Table S5**). CS and CVS values were reproducible across two cell lines (Pearson r = 0.77 and 0.58, respectively; **Fig. 6A, Fig. S6A**) indicating that these metrics reflect stable properties of RBP binding behavior rather than cell-type-specific artifacts. Conventional and unconventional RBPs, that is proteins that associate with RNA but lack known RNA-binding domains^53^, showed overlapping distributions of CS and CVS (**Fig. S6B**), suggesting that sequence discrimination, where it exists, is not restricted to proteins with canonical domain architectures.

**Figure 6:**
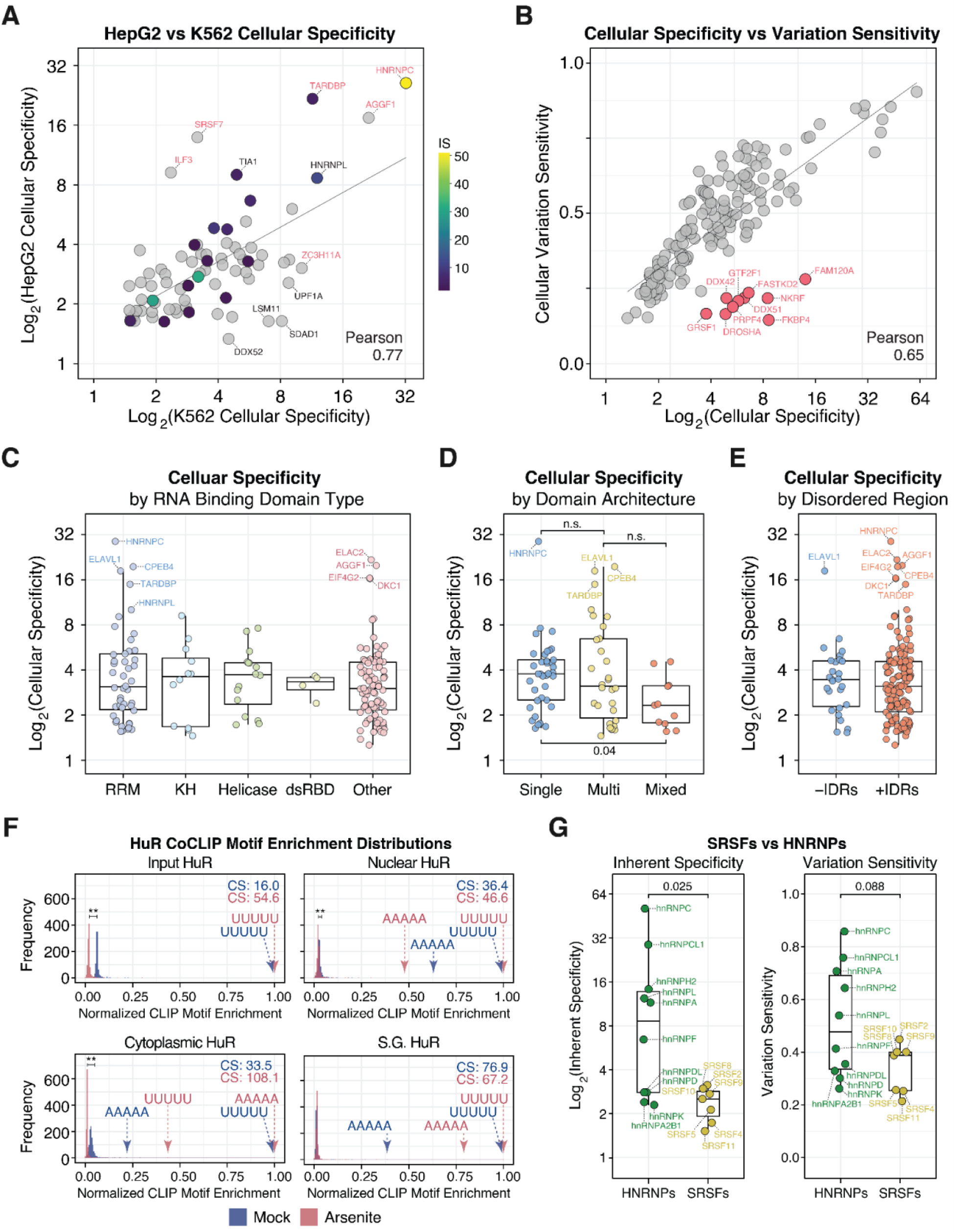
Cellular specificity and variation sensitivity analysis across an RBP repertoire. **(A)** Correlation between cellular specificity (CS) values calculated from eCLIP data in HepG2 and K562 cell lines. Pearson r correlation is shown. RBPs in red are statistical outliers; those in black deviate from the linear trendline. **(B)** Correlation between CS and cellular variation sensitivity (CVS) across different RBPs. Outlier RBPs are noted in red. **(C)** Distribution of CS by RNA-binding domain (RBD) types. Selected outliers are annotated. **(D)** Distribution of CS by domain architecture. **(E)** Distribution of CS by the presence of intrinsically disordered regions (IDRs). Statistical significance by a Wilcox-test is noted (n.s. = not significant) **(F)** 5-mer motif enrichment distributions for HuR from coCLIP experiments in total (input), nuclear, cytoplasmic, and stress granule compartments under mock or arsenite stress. CS values are indicated for each condition. Statistical significance by a two-tailed t-test is noted (**, p < 0.01). **(G)** Comparison of IS and VS values from RBNS datasets between HNRNP and SRSF protein families. A Wilcox-test result is noted.

Examining the relationship between CS and CVS across all RBPs revealed a positive correlation, consistent with *in vitro* observations (**Fig. 6B**). A notable subset of RBPs showed high cellular specificity combined with low variation sensitivity, indicating strong discrimination of a top motif alongside tolerance for single-nucleotide substitutions within it (**Fig. 6B**, highlighted in red). This combination, distinct from the high-IS, high-VS profile of HNRNPC, may reflect RBPs that recognize degenerate versions of a core motif with high selectivity against unrelated sequences. Across RBD classes, RRM, KH, helicase, and dsRBD-containing proteins showed comparable median cellular specificities and variation sensitivities, with the greatest variance among RRM containing RBPs (**Fig. 6C, Fig. S6C**). Single-domain RBPs tended toward higher variation sensitivity than those with mixed architectures (**Fig. 6D, Fig. S6D**), while the presence or absence of IDRs showed no consistent effect on either metric (**Fig. 6E, Fig. S6E**).

Our colocalization CLIP data^47^ for HuR further illustrates that cellular specificity is not a fixed property of an RBP. HuR showed higher cellular specificity in cytoplasmic and stress granule fractions relative to nuclear fractions (**Fig. 6F**), consistent with compartment-specific changes in target availability and RBP competition. At the family level, HNRNPs showed significantly higher IS values than SRSFs (**Fig. 6G**), whose lower specificity is consistent with their role in recognizing exonic and intronic splicing enhancers, in contrast to the more selective silencer-element targeting characteristic of HNRNPs^54,55^. Together, these analyses show that IS and VS capture meaningful variation in RNA binding behavior across the RBP repertoire, from individual proteins to protein families. IS, VS, and cellular specificity values for all RBPs analyzed here are provided in **Table S5**, and can be computed for any RBNS or CLIP dataset using the RBPSpecificity R package, available on GitHub.

## Discussion

In this work, we describe two metrics, inherent specificity (IS) and variation sensitivity (VS), to quantify and compare RNA-binding specificity across RBPs. IS assesses how strongly an RBP discriminates its top binding motif from the full landscape of potential targets, while VS evaluates its sensitivity to single nucleotide changes within that motif. Although both metrics focus on the strongest binding motif, IS captures global discrimination across sequence space, while VS captures local discrimination within the immediate sequence neighborhood of the optimal motif. IS and VS reduce complex motif enrichment distributions into standardized, protein-independent quantities that enable systematic cross-RBP ranking and serve as inputs for competitive binding models, analogous to how *K_d_* enables comparison of binding affinities. Applying these metrics across 26 single-domain RBPs *in vitro*, we find that IS and VS are inter-correlated and that RBP rankings are preserved across motif lengths from 4 to 7 nucleotides, indicating that both metrics reflect stable properties of each RBD rather than artifacts of the sequence length used to measure them.

Comparing *in vitro* and cellular measurements across 28 RBPs, we observe that correspondence between inherent and cellular specificity depends on whether binding is driven by sequence or local RNA structure. RRM-swap experiments between HNRNPC and RBM25 demonstrate that specificity can be transferred between protein contexts and quantitatively tracked using CS and CVS. Finally, mathematical modeling and competition CLIP experiments reveal that low-specificity RBPs can focus the binding of high-specificity partners by occupying non-target sites, an emergent competitive property not predictable from affinity alone. To encourage broad application of these metrics, we have implemented IS and VS calculations in the RBPSpecificity R package and competitive binding simulations in RBPEqBind. Both are available on GitHub, making them accessible to any lab working with RBNS or CLIP data.

The question of what constitutes binding specificity can be framed at multiple levels of resolution. At the atomic level, specificity arises from direct contacts between protein side chains and RNA nucleobases, whereas backbone contacts, which are sequence-independent, are generally considered non-specific^12,13^. IS and VS operate at the level of the full domain or protein, characterizing how binding is distributed across sequence space rather than describing individual contacts. One practical consequence is that the k-mer length at which IS stabilizes likely reflects the structural contact footprint of the domain, typically 3-4 nucleotides for RRMs and KH domains, beyond which additional sequence space is added without additional discriminating contacts. Rank order across RBPs is preserved regardless of k-mer length, making cross-RBP comparisons robust to this choice.

To test whether IS and VS can track the transfer of specificity between protein contexts, we performed RRM-swap experiments between HNRNPC and RBM25, two single-RRM proteins at opposite ends of the specificity spectrum. Substituting the high-specificity HNRNPC RRM into RBM25 shifted RBM25 binding toward UUUUU motifs, increasing both CS and CVS, consistent with the introduction of base-specific contacts characteristic of the HNRNPC RRM^56^. The reciprocal swap had little effect, consistent with HNRNPC sequence selectivity being distributed across multiple structural elements beyond its RRM. In addition to the RRM, HNRNPC contains a basic region that contributes sequence-independent RNA affinity^57,58^, functions as a homotetramer that spaces binding cooperatively along poly-U tracts^34,48,50^, and is embedded in the Large Assembly of Splicing Regulators (LASR) complex^59^. That CS and CVS detected the partial transfer of specificity in the RBM25 direction while reflecting the resistance of HNRNPC to modification confirms that the metrics are responsive to changes in specificity where it is RRM-encoded, and informative about where it is not.

Cellular specificity also appears to vary based on the environment in which an RBP operates. For example, our HuR coCLIP analysis show that cellular specificity shifts with subcellular localization and stress, with cytoplasmic and stress granule-associated HuR displaying higher specificity than nuclear fractions, suggesting the effects of compartment-specific differences in target availability and RBP competition. At the family level, SRSFs show lower IS and VS values than HNRNPs, a separation that aligns with their antagonistic roles as splicing activators and repressors respectively^54,55^ and suggests that IS and VS reflect meaningful differences in how RBP families engage their targets. Within this broader pattern, a subset of RBPs in Figure 6B shows moderate cellular specificity paired with low variation sensitivity, suggesting that global discrimination across sequence space and local sensitivity to substitutions within the top motif can be partially decoupled.

How specificity profiles translate into target selection in a cellular environment where RBPs compete for overlapping sequence space remains an open question. Prior quantitative modeling established that as binding sites in the transcriptome exceed the available pool of regulators, occupancy at any given site is shaped by competition across the full sequence space an RBP encounters^60^. In this view, non-”productive” binding could shape the availability of binding at preferred sites, analogous to the sponging and buffering effects described for miRNAs^60,61^. Our competition modeling and cellular competition CLIP experiments provide experimental support for this principle where low-specificity RBPs enhance the target discrimination of high-specificity partners by occupying non-preferred sites. Most CLIP-detected binding events may reflect this kind of distributed occupancy rather than direct regulatory consequence at each site. IS and VS provide a quantitative handle on this question by characterizing where in sequence space each RBP concentrates its binding.

Several directions remain open. IDR-containing RBPs present an immediate challenge as their RNA contacts are multivalent^21^, making it unclear whether an affinity distribution derived from CLIP or *in vitro* binding data reflects intrinsic sequence preference or proximity-driven association. More broadly, there is growing appreciation that RNA itself can act as a regulatory player, organizing protein complexes and impacting their activity through structural and sequence features rather than through classical RBD-mediated recognition^13^. In this view, low apparent specificity need not reflect promiscuity but may instead reflect a mode of engagement in which RNA recognition is an emergent property of the complex rather than any individual component. Furthermore, sequence specificity can emerge from the assembly of multi-protein complexes whose individual components show little selectivity on their own, as has recently been demonstrated^62,63^. How IS and VS behave for IDR-containing proteins and larger assemblies is an open question, particularly as higher-resolution methods for mapping IDR-RNA contacts and complex-level binding preferences become available. Recent large-scale *in vitro* efforts have additionally expanded the number of RBPs with characterized motif data across eukaryotes^64^, and these datasets are compatible with IS and VS calculation using RBPSpecificity, providing a ready starting point for applying these metrics at repertoire scale as matched cellular data become available.

Lastly, these metrics provide a rational basis for protein engineering efforts aimed at designing RBPs with targeted binding properties^65–69^. Cellular specificity is also shaped by post-translational modifications and subcellular localization, which alter either RNA contact geometry or target availability in ways that are not captured by *in vitro* measurements alone^70,71^. Emerging approaches for subcellular RNA profiling and proximity-based RBP-target mapping will be essential for characterizing these context-dependent shifts systematically^47,72–74^.

### Limitations of Study

Our study is not without limitations. We treat CLIP peaks as evidence of RBP presence at a given site without making claims about binding stoichiometry or functional consequence, which is the same assumption made by most motif-finding approaches applied to CLIP data. We also acknowledge that motif enrichment derived from CLIP experiments are not direct measures of cellular binding affinities. Measuring true cellular affinities across all possible k-mers for a given RBP remains a significant challenge; we therefore adopt the formal assumption that enrichment-based rankings approximate the underlying affinity landscape in cellular settings, motivated by the strong correlation between RBNS motif enrichment and relative *in vitro* affinity ^28^. In addition, IS and VS require sufficient sampling of sequence space to generate a meaningful motif enrichment and relative affinity distribution, and for RBPs with sparse CLIP peaks the resulting metrics will be poorly constrained.

These caveats apply to unconventional RBPs such as proteins identified through RNA interactome capture that lack annotated RBDs and show little or no sequence specificity in orthogonal assays^53^. For such proteins, low IS values may reflect genuine sequence promiscuity, technical limitations of crosslinking-based methods, or indirect RNA association through protein complexes rather than intrinsic binding. Applying IS and VS systematically to unconventional RBPs is nonetheless worth pursuing. Proteins that lack sequence preference should produce predictably low and flat affinity distributions, and deviations from that baseline, whether across conditions, cellular states, or complex partners, would themselves be informative. More broadly, resolving the questions raised here will require advances in crosslinking chemistry, single-molecule binding assays, and methods that capture RBP binding within intact complexes rather than in isolation. As these approaches mature, IS and VS provide insights for interpreting what they reveal about the full spectrum of RNA recognition strategies operating in cells.

## Data and code availability

All original code for analysis and data visualization is available on GitHub at https://github.com/S00NYI/BITS_Specificity. To facilitate broader application of our metrics, we have developed an R package that enables straightforward calculation of IS and VS values from sequencing data. It is available at https://github.com/LunaRNALab/RBPSpecificity. In addition, we have developed an R package for RNA-RBP binding simulations, which is available at https://github.com/LunaRNALab/RBPEqBind. All the sequencing data generated by this study have been deposited in the NCBI Gene Expression Omnibus (GEO) database under accession number GSE291358 and GSE326416.

## Acknowledgments

We are grateful to William Merrick, Maria Hatzoglou, and Boaz Tirosh for their helpful discussion and feedback. We further thank Frank Tedeschi, Thomas J. Sweet, and Daniel Dominguez for feedback and a critical reading of this manuscript. pTRIPZ shNS was a gift from Sandra Demaria. This work was supported in part by the NIH (T32 GM152319 to S.Y., R35 GM118088 to E.J., R35 GM154771 to J.M.L.), the Case Comprehensive Cancer Center (P30 CA043703 to J.M.L.), the American Cancer Society (IRG-16-186-21 to J.M.L.), as well as startup funds (to J.M.L.) from the Department of Biochemistry at the Case Western Reserve University School of Medicine.

## Author Contributions

Conceptualization, S.Y., X.Y., E.J.; Investigation, S.Y., S.S.S., R.K., V.K., J.M.L; Writing – Original Draft, S.Y. and J.M.L; Writing – Review & Editing, S.Y., E.J., J.M.L.; Funding Acquisition, E.J. and J.M.L.; Supervision, E.J. and J.M.L.

## Declaration of Interests

E.J. is a member of the scientific advisory board of Eclipsebio. The other authors declare no competing interests.

## Materials and Methods

### Plasmid DNA Construction

To generate stable cell lines expressing the wild-type and the mutant HNRNPC and RBM25, we cloned each RBP construct into an enhanced Piggybac (ePB) doxycycline-inducible expression vector^75^. The wild-type and the mutant HNRNPC construct was prepared as a custom gBlock from Integrated DNA Technologies. The wild-type RBM25 construct was prepared as cDNA reverse transcribed using total RNA extracted from HEK293 cells. For the mutant RBM25 construct, the DNA segments encoding for RBM25 RRM was replaced with HNRNPC RRM through overlap PCR. Finally, a V5 epitope tag^76^ was appended to the N-terminus of all constructs. Synthesized constructs were cloned between BamHI and NotI sites in ePB vectors. For the endogenous HNRNPC knockdown, short hairpin RNA (shRNA) sequences were cloned into the pTRIPZ vector (Addgene cat# 127696)^77^ between the *Xho*I and *Eco*RI restriction sites. All primers and shRNA sequences used to generate these constructs are listed in Table S6.

### Cell Culture and Stable Cell Line Generation

HEK293 cells^78^ were maintained at 37°C and 5% CO_2_ in Dulbecco’s Modified Eagle Medium (DMEM, Fisher Scientific, cat. #11995065) supplemented with 0.1 mM nonessential amino acids (NEAA, Fisher Scientific, cat. #11140076) and 5% hyclone fetal bovine serum (FBS, ATLAS Biologicals, cat. #F-0500-D, Lot # F05D22D1). All cell lines tested negative for mycoplasma contamination.

Stable cell lines were generated by transfecting 1μg of ePB plasmid encoding each RBP construct with 1μg of pTransposase plasmid^75^ using Lipofectamine 2000 (Invitrogen, cat. #11668027). Cells underwent selection with 1μg/ml Puromycin (Sigma, cat. #P8833-25MG) to eliminate non-transfected cells. Doxycycline (Sigma, cat. #D9891-1G) was used at 1μg/ml to induce RBP expression.

Expression of the RBPs and their localizations were confirmed through western blot analysis and immunofluorescence imaging. For western blot, HEK293 cell lysates were prepared in 1x PXL lysis buffer supplemented with complete protease inhibitors (Roche, cat. #11873580001). All lysates were prepared from equal cell numbers per condition. Protein concentrations were determined by Bradford assay (Biorad), and 20μg total protein per sample was run on NuPAGE gels (Life Technologies) and transferred to fluorescence-compatible nitrocellulose membranes (Millipore). Membranes were blocked in Odyssey PBS-based buffer (LI-COR) for 1 hr, then primary antibodies were added for an overnight incubation at 4°C. Antibodies used for western blotting were: HNRNPC (ThermoFisher Scientific, cat. #PA5-22280; 1:1000), RBM25 (ThermoFisher Scientific, cat. #PA-90511, 1:1000), rabbit V5-tag (Cell Signaling Technology, cat. #13202S, 1:1000), mouse V5-tag (Cell Signaling Technology, cat. #80076S, 1:1000), and Beta-Actin (Sigma, cat. #A5441, 1:5000). After 3 washes in 1X PBS/0.05% Tween-20, membranes were incubated fluorescent secondary antibodies (LI-COR, 1:25,000) for 1 hr at room temperature. Membranes were washed 3 times in 1X PBS/0.05% Tween-20, rinsed in 1X PBS, and visualized on the Odyssey Imaging system (LI-COR).

For immunofluorescence, cells were plated either on glass coverslips or black-walled clear bottom 96-well plates (Corning, Cat# 3904) that were coated with poly-L-lysine. Upon harvest, cells were fixed with 4% paraformaldehyde (PFA) for 10 minutes. PFA was then removed, and cells were stored at 4°C in PBS containing 1% FBS until processing. Cells were washed in PBS containing 0.1% Tween-20 (PBST), permeabilized with PBS containing 0.1% Triton X-100 for 10 min at room temperature, and blocked for 1h at room temperature with a blocking solution of 5% BSA in PBST. Cells were stained with primary antibodies overnight at 4°C. Antibodies used for immunofluorescence were: HNRNPC (ThermoFisher Scientific, cat. #PA5-22280; 1:1000), RBM25 (ThermoFisher Scientific, cat. #PA-90511, 1:1000), rabbit V5-tag (Cell Signaling Technology, cat. #13202S, 1:1000), and mouse V5-tag (Cell Signaling Technology, cat. #80076S, 1:1000). After primary antibody incubation, cells were washed and stained with goat anti-rabbit IRDye 680RD antibody (LICOR, cat. #925-68971, 1:1000) and goat anti-mouse IRDye 800CW antibody (LICOR, cat. #925-32210, 1:1000). Nuclei were counterstained with DAPI (ThermoFisher Scientific cat. #D1306, RRID:AB_2629482) at 1 µg/ml, for 5 minutes prior to imaging. Fluorescent images were obtained on a Keyence BZ-X710 microscope or on a Cytation 7 (Agilent).

### Endogenous HNRNPC Knockdown

To knock down endogenous hnRNPC, doxycycline-inducible hnRNPC wild-type and mutant overexpression cell lines were transfected with shRNA constructs (pTRIPZ-shHNRNPC or pTRIPZ-NS control). Because both the overexpression system and shRNA are doxycycline-inducible, parallel experiments were performed in HEK293 cells to assess knockdown of endogenous hnRNPC. shRNA expression was induced with doxycycline (1 µg/mL) for 48 h prior to harvesting for Western blot analysis. The shRNAs used targeted the HNRNPC 3′UTR, thereby reducing endogenous hnRNPC without affecting exogenous cDNA-derived expression.

### RBP CLIP

CLIP from crosslinked HEK293 cell pellets using 5µg per sample of mouse V5-tag antibody (Cell Signaling Technology, cat. #80076S) or 2µg per sample of mouse monoclonal HNRNPC antibody (Santa Cruz Biotechnology,, cat. #sc-32308) was performed following previous work^47,79,80^ with modifications. Detailed experimental procedures are outlined below.

### Tissue culture and crosslinking

Cells were plated onto 10 or 15 cm dishes, and expression was induced for 2 days with 1μg/ml of doxycycline. The culture media was replaced with ice-cold PBS and cells were immediately irradiated under 254nm UV light on ice, once for 400 mJ/cm^2^ and once again for 200 mJ/cm^2^ using a Stratalinker 2400 (Agilent Genomics). These energies were based on our previous work^47,79,80^. Cells were scraped into tubes and pellets were stored at −80°C until downstream preparation.

### Bead preparation

For V5 antibody pull downs, protein G Dynabeads (Invitrogen, cat. #10004D) were washed 3x with and resuspended in antibody binding (AB) buffer (AB: PBS, 0.02% Tween-20). Beads were incubated with 5μg of Anti-V5 [IPI-SV5-Pk1] antibody^81^ from Institute for Protein Innovation (Addgene antibody # 218107) per 25μL beads for 30 min at room temperature or overnight at 4°C. Prior to IP, beads were washed in 1x PXL lysis buffer (1x PXL: 1x PBS tissue culture grade without magnesium or calcium, 0.1% SDS, 0.5% Sodium-deoxycholate, 0.5% NP-40, with protease inhibitors).

### V5 IP and on-bead enzymatic steps

Lysates from crosslinked cells were prepared by adding 1ml of 1x PXL lysis buffer + quenchers + protease inhibitors (Roche, cat. #11873580001) and triturating to disrupt cells. Lysates were treated with 10μL DNase (RQ1, Promega) and placed on ice for 10 minutes. Lysates were then treated with RNase I (Thermofisher, cat. #EN0602), first diluted to the indicated concentration by volume (1:100 for over digest or ‘OD’, or 1:1,000 for Low RNase) in 1x PXL and then added at 10μl per mL of lysate. RNase I concentrations were empirically determined for optimal isolation of RNA-RBP complexes with sizes suitable for cloning (Moore et al. 2014). Lysates underwent thermomixing for 5 minutes at 37°C at 1100rpm before being spun at 4°C on max speed of a table-top microcentrifuge for 10 minutes. All subsequent steps were done on ice or at 4°C unless otherwise indicated. Supernatants, along with any lipid layer, were harvested and mixed with PXL-equilibrated V5 antibody-bound beads for immunoprecipitation. Samples were nutated with beads at 4°C for 2 hours. Beads were washed sequentially twice each with ice-cold 1x PXL, 5x PXL (same as 1x but using 5x PBS), and 1x PNK buffer (50mM Tris-HCl, pH 7.5, 10mM MgCl2, 0.5% NP-40).

To prepare RNA 3’ ends for linker ligation, IPs were treated with alkaline phosphatase. Beads were resuspended in 40μL containing 1x dephosphorylation buffer, 3U of CIAP (Roche), RNasin inhibitor (Promega), and thermomixed for 20 minutes at 37°C, shaking at 1100rpm for 15 s every 2 min. Samples were washed as above sequentially in ice-cold 1x PNK, 1x PNK plus 20mM EGTA, and twice with 1x PNK.

Radiolabeled linkers were prepared with a poly-nucleotide kinase (PNK) reaction consisting of 100pmol of a 3’ inverted ddT blocked L32 RNA linker, 0.5μL of 32P-gamma-ATP (Revvity, cat. #NEG035C005MC), 0.5µl 1x T4 PNK buffer, 0.25μL T4 PNK and 0.3μl RNasin inhibitor in a total volume of 5µl per sample. Radiolabeled linker was prepared for 10 ligation reactions in a volume of 50µl. Linkers were incubated for 20 min at 37°C, after which 2µl of 10mM ATP was added, and the reaction was incubated at 37°C for an additional 5 minutes. Linker was purified by passing through a G-25 column (GE Healthcare, cat. # 27-5325-01) and stored at −20°C until use.

Linker ligation at 3’ ends was set up per sample after washes following alkaline phosphatase treatment by preparing a T4 RNA ligase 1 (NEB, cat. #M0204S) reaction in 40μL following the manufacturers’ instructions with 100pmol of radiolabeled L32 RNA linker. Samples were incubated overnight at 16°C, shaking at 1100rpm for 15 s every 4 min. The next day, beads were washed twice each with 1x PXL and 5x PXL, before being equilibrated in 1x PXL.

### RNA-RBP complex isolation and autoradiography

All beads were subsequently washed twice each with 1x PXL, 5x PXL, and 1x PNK. Protein was eluted off the beads by incubating with 30µl of 1X LDS loading buffer (Invitrogen) without reducing agent for 10 min at 70°C, shaking at 1100rpm. Supernatants were run on Novex NuPAGE 8% Bis-Tris gels (Invitrogen, cat. #WG1001BOX) in SDS-MOPS buffer at 4°C. Radiolabeled protein RNA complexes were transferred to BA85 nitrocellulose (Cytiva Amersham, cat. #45-004-007). After transfer, the membrane was rinsed with RNase-free PBS, and exposed to Biomax MR film (Kodak) at −80°C from 3 hr to up to 3 days. Alternatively, membranes were placed onto phosphor screens (GE cat. # BAS-IP VS 2025), and imaged with a Typhoon Scanner (GE Amersham).

Nitrocellulose membranes were aligned with the exposed film and regions of the membrane from low RNase IP lanes were excised corresponding to signal intensities for RNA-RBP complexes between 42-72kDa for HNRNPC and 160-180kDa for RBM25. The 10-30kDa above the RBP size region (32kDa for HNRNPC, 150kDa for RBM25) was chosen to confine the CLIP-seq analysis to RNA target fragments of 30-90nt nucleotides (assuming 3nt/kDa) to avoid confounding variables from higher molecular weight complexes. RNA was liberated from membrane fragments using 200µL of a 4mg/ml proteinase K solution (Roche, cat. #3115828001) diluted in PK buffer (100mM Tris-HCl, pH 7.5, 50mM NaCl, 1mM EDTA, 0.2% SDS) and incubated for 60 min at 50°C, shaking at 1100rpm for 15 s every 2 min. RNA fragments underwent acid phenol:chloroform extraction using and were precipitated overnight at −80°C. RNA was pelleted by spinning at max speed (> 13,000rpm) in a table top centrifuge at 4°C, and washed twice with 75% ethanol. Following the drying of the RNA at the bench, the pellet was dissolved in 8µL RNase-free water.

### RBP footprint library generation and sequencing

CLIP footprints were reverse-transcribed using the Br-dU incorporation and bead-capture strategy described previously^82,83^. Indexed reverse transcription (RT) primers were used (listed in Table S6), allowing multiplexing of 22 samples per Nextseq 500 run. cDNA was circularized with CircLigase (Epicentre), pooled and then amplified with PCR primers with Illumina sequencing adapters as described previously^82,83^. Amplification was tracked with SYBR green (Life Technologies) on the Quantstudio real-time PCR machine (Thermo), and reactions were stopped once signal reached 100k relative fluorescence units (r.f.u.). Products were purified with Ampure XP beads (Beckman) and quantified via Qubit assay (ThermoFisher) and/or Tapestation system (Agilent). Multiplexed samples were run on the Illumina NExtSeq Mid-Output with 75 base pair single-end reads.

### Bioinformatics Analysis

Scripts and processed datasets used for the analysis and generation of figures in this article are available on GitHub (https://github.com/S00NYI/BITS_Specificity). For the CLIP analysis, our single command line CLIP analysis tool ‘CLIPittyClip’ can be found on our GitHub repository https://github.com/LunaRNALab/CLIPittyClip. In addition, custom R packages ‘RBPSpecificity’ for motif enrichment analysis and ‘RBPEqBind’ for RNA-RBP binding simulation are available on their respective GitHub repositories at https://github.com/LunaRNALab/RBPSpecificity and https://github.com/LunaRNALab/RBPEqBind.

### RBNS data collection and processing

RBNS datasets were collected from the ENCODE portal^40^ by searching “RNA Bind-n-Seq” in “Experiment search.” RBNS motif enrichment measurements for each RBP are performed at several different RBP concentrations and analyzed for different *K*-mer lengths. For each *K*-mer enrichment data, the RBP concentration that generated the highest R-value was selected as the representative sample and used throughout the analysis. The RBNS dataset that used only a single purified RNA binding domain as well as RBPs’ structural context preferences were manually curated based on previous literature^29^. A list of the RBNS accession numbers is provided in Table S1. The collected RBNS *R* scores were then feature-scaled to a scale of 1-to-Euler’s constant (*e*), followed by natural log transformation to fit the data into a scale of 0-to-1^42^. All normalized relative affinity values datasets used and associated IS/VS values are provided in Table S2 and S3, respectively.

### PWM and sequence logo generation

From RBNS datasets, sequence variants with RBNS R scores two standard deviations above the average R score were used as input for the *R* program *consensusMatrix* under the *Biostrings* package^84^. The output nucleotide frequency table was then used as an input for the R program *PWM* under the *Biostrings package* to generate the PWM for each tested RBP. To generate a sequence logo, the generated PWM was used as an input for *R* program *seqLogo*^85^ with an ‘RNA’ option to generate the sequence logo. To calculate the predicted binding scores based on the PWM, the sum of the weights for a nucleotide at each position was calculated for all possible 5-mers. The calculated binding scores were then normalized as described before.

### IS and VS calculation

Inherent specificity measures the degree by which an RBP differentiates the strongest binding *K*-mer motif from the rest of the *K*-mers. Inherent specificity is calculated by taking the ratio of the top affinity over the median affinity of the affinity distribution. Numerically, inherent specificity is calculated as:

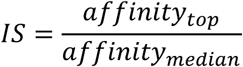

Variation sensitivity measures the degree of changes in binding affinity for an RBP when a single nucleotide is changed in its strongest binding *K*-mer motif. Variation sensitivity is calculated by taking the average changes in affinity between the strongest binding motif and the motif with one nucleotide changed from the strongest binding motif. Numerically, variation sensitivity is calculated as:

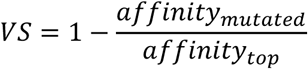

For the cellular specificity and cellular variation sensitivity calculations, eCLIP motif enrichments derived from the ENCODE peak files were used instead of RBNS affinity measurements. See CLIP Dataset Processing and Analysis section of the Methods for more detail.

### CLIP Dataset Processing and Analysis

eCLIP datasets were collected from the ENCODE portal^40^ by searching “eCLIP” in “Experiment search.” For each RBP, merged bed narrowPeak files were downloaded. For the HNRNPC/RBM25 CLIP, HNRNPC competition CLIP, and HuR coCLIP datasets, the raw sequencing data was processed using the *CLIPittyClip* pipeline as described before^47^ to generate the peak matrices. All identified peaks were annotated by genomic region using GENCODE annotations. Using the peak coordinates, 5-mer motif enrichment for each CLIP experiment was calculated as local background corrected 5-mer motif frequency near peaks using the motifEnrichment function in the RBPSpecificity R package. For each peak, the local background was defined as a region of the same length as the peak with at least 500 nucleotides but no more than 1000 nucleotides up or downstream from the peak. The background 5-mer motif frequency profile was bootstrapped 100 times to normalize for any selection biases. The averaged background motif frequency was then subtracted from the motif frequency at peak coordinates to generate the motif enrichment profile. For the peak motif enrichment, following nucleotide extension or trimming was performed for each CLIP dataset: 25 nucleotides extension at the 5’-end for eCLIP peaks; 15 nucleotides extension at the 5’-end and 15 nucleotides trimming at the 3’-end for HNRNPC and RBM25 CLIP peaks; 10 nucleotides extension at the 5’-end and 5 nucleotides trimming at the 3’-end for Hur coCLIP peaks. For more detail on the specific implementation and usage of the R package for motif enrichment analysis, please refer to GitHub repositories (https://github.com/S00NYI/BITS_Specificity and https://github.com/LunaRNALab/RBPSpecificity).

For the per-peak tag enrichment analysis in the competition CLIP experiment, HNRNPC peaks identified across all conditions were used as a common coordinate set. For each peak, sequence depth normalized tag counts were calculated under each condition (no overexpression as *null*, WT RBM25 overexpression, and Mut RBM25 overexpression). Per-peak tag enrichment was then calculated as the ratio of depth-normalized tag counts upon RBM25 overexpression to depth-normalized tag counts in the *null* condition, separately for WT and Mut RBM25. Statistical significance between the WT and Mut RBM25 overexpression enrichment distributions was assessed by a Kolmogorov-Smirnov test.

### RNA-RBP binding simulation

RNA-RBP binding simulations in different conditions were performed using the *RBPEqBind* R package. Because motif enrichment distributions do not directly measure Kd values but correlate strongly with measured relative affinities in RBNS^28^, we used normalized relative affinity distributions derived from enrichment rankings as a proxy for the binding affinity landscape of each simulated RBP. The four hypothetical RBPs with different 5-mer IS and VS combinations were created by curating the normalized relative affinity distributions. Four different relative affinity distributions for the 1,024 5-mer motif affinities were first created so that they follow a normal distribution. For simplicity, all four RBPs were set to bind ‘UUUUU’ with the highest affinity. Then, for RBPs with high IS, 5-mer motifs excluding the top binding motif were shifted to a lower affinity regime. Then, for RBPs with low VS, the 5-mer motifs with single nucleotide mutations from the top binding motif were placed near the top binding motif. For the RBPs with high VS, the same set of 5-mer motifs was placed at the lowest end of the affinity distribution. Finally, the affinities for motifs that are neither the top binding motif nor motifs with single nucleotide changes from the top binding motifs were shuffled and curated to give each RBP a unique binding affinity landscape. The *in silico K_d_* towards the top motif for the RBPs was all set to 0.01 nM and *K_d_* values for other motifs were scaled according to their relative positions in the normalized affinity distribution.

For a given RNA sequence of length *l* that interacts with two or more RBPs, there exists *l* − *(k*−1) possible *K*-mer binding sites. For each binding site on the RNA, a kinetic reaction scheme can be constructed where *P_i_* is an *i*^th^ out of *N* RBP interacting with the binding site *R* on the given RNA:

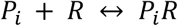

Then, at equilibrium:

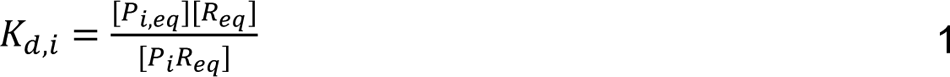

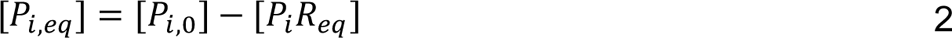

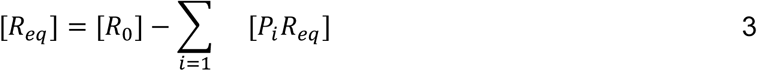

where *K_d_*_,i_ is the dissociation constant for *P_i_* and [*P_0,i_*] and [*R_0_*] are the initial concentrations of the RBPs and RNA, respectively. Combining equations 1 and 2, and solving for the equilibrium concentration of the *P_i_R* complex yields:

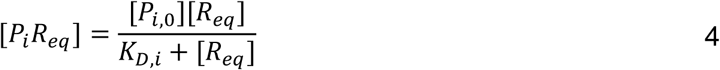

Combining equations 3 and 4 yields an equation in terms of initial conditions and the equilibrium concentration of the binding site:

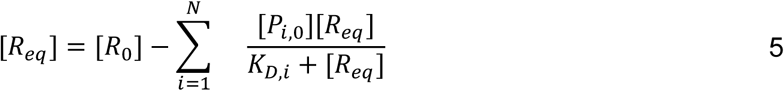

For the system of *N*=4, solving equation 5 in terms of [*R_eq_*] yields a 5th-degree polynomial, which can be solved with the boundary condition:

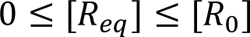

From the [*R_eq_*], the equilibrium concentration of the *P_i_R* complex can be determined with equation 4.

For each binding site, effective initial RNA concentration needs to be determined as a given binding site may exist more than once in the given RNA sequence. Thus, the effective initial concentration of a binding site [*R_0,e_*] is calculated as *n* x [*R_0_*], where *n* is the number of the binding site motif appearances in the given RNA sequence.

Dividing the equilibrium *P_i_R* complex concentration [*P_i_R*] by the initial concentration of the RNA [*R_0_*] yields a probability of the RNA binding site occupied by the protein, *p_site_*. Using *p_site_* for all possible binding sites on a given RNA, per position probability, *p_pos_*, can be calculated by taking the average of the overlapping *p_site_*for each position (up to *k* overlap). Then, for any given *K*-mer motif within the RNA sequence, average probability can be calculated by taking the average of the *k p_pos_* that span the motif.

For RBP enrichments across the target RNA, the proportion of each RBP distributed across the RNA was calculated by normalizing the equilibrium *P_i_R* complex concentration [*P_i_R*] by the sum of the equilibrium concentration for each RBP. Then, the RBP occupancy was calculated by calculating the proportion of the RBP at a given position compared to the initial concentration.

### RRM Structure Prediction and Comparison

Amino acid sequences of RRMs of RBFOX3, HNRNPC, RBFOX2, HNRNPCL1, RALYL, RBM41, TRA2A, PABPN1L, RBM24, RBM6. SRSF2, SRSF8, SRSF10, RBM25, and SRSF11 were acquired through the UniProt database^86^. The full length amino acid sequence was used as input for AlphaFold protein structure prediction program^87^ and UCSF ChimeraX software was used to visualize the structures^88^. Amino acid sequence alignment was performed through MUSCLE alignment software^89^.

## Statistical Analysis

All statistical analyses were performed in R using the *stats* package.

## Notes

https://www.ncbi.nlm.nih.gov/geo/query/acc.cgi?acc=GSE291358

https://github.com/S00NYI/BITS_Specificity

https://github.com/S00NYI/RBPSpecificity

https://github.com/S00NYI/RBPEqBind

https://www.ncbi.nlm.nih.gov/geo/query/acc.cgi?acc=GSE326416

### Summary of Updates

Abstract revised, author contact info updated

