## Supplementary Figures for "Inherent Specificity and Variation Sensitivity as Quantitative Metrics for RBP Binding"

**Supplemental Items:**
Figure S1: Inherent specificity and variation sensitivity across k-mers and RRM structural/sequence alignments of single-RRM proteins. (related to Figures 1 and 2)
Figure S2: RBP CLIP optimization and autoradiograms. (related to Figures 3 and 5)
Figure S3: Characterization of WT and mutant HNRNPC CLIP with endogenous HNRNPC knockdown. (related to Figures 3 and 5)
Figure S4: *In silico* simulation results of RBP competitive binding on target RNA. (related to Figure 4)
Figure S5: *In silico* simulation and peak analysis results of Competition CLIP (related to Figure 5)
Figure S6: Cellular variation sensitivity analysis across an RBP repertoire. (related to Figure 6)

**Supplemental Table:**
Table S1: List of ENCODE accession IDs of RBPs examined (related to Figures 1-3, 6, S1, S2, S3, and S5)
Table S2: Normalized Relative Affinities for 4 to 7-mers RBNS dataset (related to Figures 1-2, and S1)
Table S3: IS and VS calculated for 4 to 7-mers RBNS dataset (related to Figures 1-2, and S1)
Table S4: Normalized Relative Affinities for Model RBPs and Motif Enrichments for WT HNRNPC, and WT/Mut RBM25
Table S5: 5-mer IS/VS and CS/CVS values for ENCODE eCLIP RBPs (related to Figure 6 and S5)
Table S6: Oligonucleotides (related to Figures 3 and 5)

### Figure S1

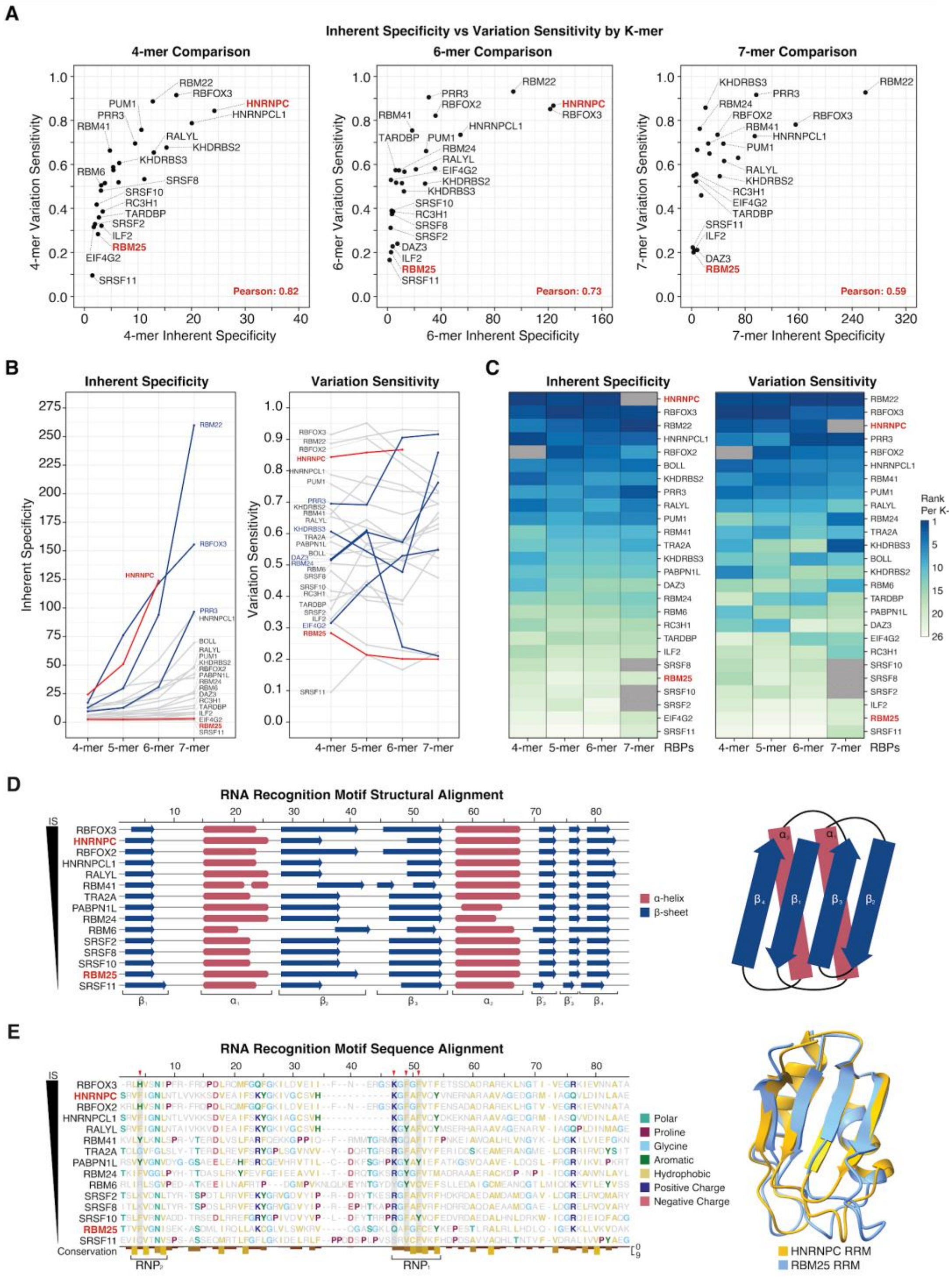


**Figure S1: Inherent specificity and variation sensitivity across k-mers and RRM structural/sequence alignments of single-RRM proteins. (A)** Correlation between IS and VS for 4-mer, 6-mer, and 7-mer sequences. Pearson correlation coefficients are indicated in red. **(B)** Line graphs showing the changes in IS and VS values across 4 to 7-mer. RBPs of interest (HNRNPC and RBM25) are noted in red. **(C)** Heatmaps showing the rank of Inherent Specificity

(IS) and Variation Sensitivity (VS) for 26 single-RRM RBPs across different k-mers. **(D)** Structural alignment of RRMs from the 26 RBPs, ordered by descending IS. Secondary structure elements (α-helices in red, β-sheets in blue) were predicted using AlphaFold. A schematic of the canonical RRM fold is shown on the right. **(E)** Protein sequence alignment of the 26 RRMs with amino acids colored by chemical properties. RNP1 and RNP2 motifs are indicated at the bottom. The right panel shows the structural superimposition of the AlphaFold-predicted HNRNPC (yellow) and RBM25 (blue) RRMs.

Figure S2

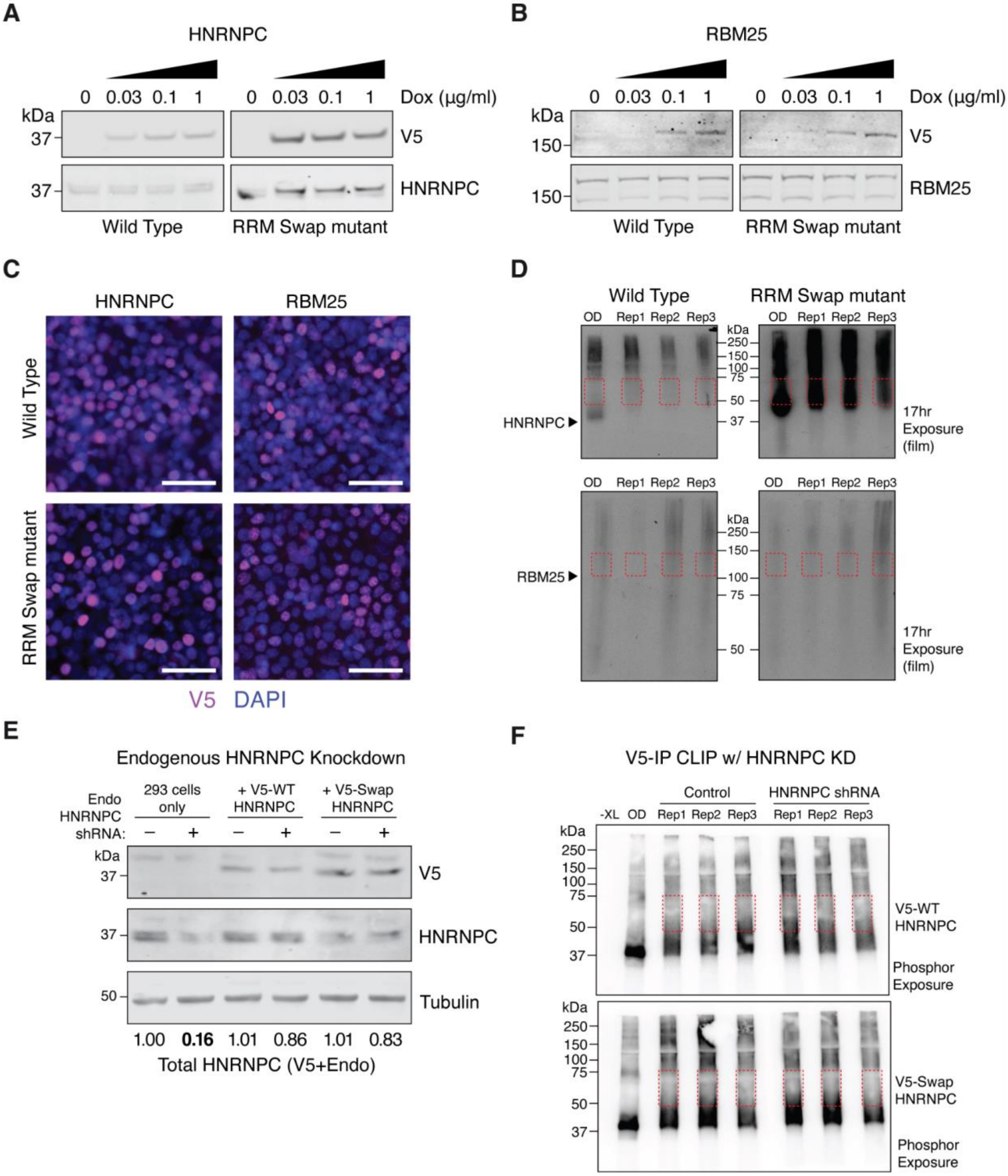


**Figure S2: HNRNPC and RBM25 CLIP optimization and autoradiograms. (A)** Western blot analysis of V5-tagged wild-type and the RRM-swap mutant HNRNPC. The blot was probed with both anti-V5 antibody and anti-HNRNPC antibody to confirm exogenous and total expression, respectively. The amount of doxycycline used to induce exogenous expression is shown. **(B)**

same as **(A)** but for RBM25. **(C)** Immunofluorescence staining of HEK293 cells stably expressing the wild-type and the mutant HNRNPC and RBM25. 1ug/mL of Doxycycline was used to induce RBP expression. Anti-V5 antibody was used to confirm expression and localization of the RBPs. DAPI used as DNA counterstain. Images were captured at 20x magnification. Scale bars = 10μm. **(D)** Autoradiogram of HNRNPC (top) and RBM25 (bottom) CLIP after V5 antibody IP. OD: RNAse overdigest. Excised regions for sequencing indicated with red boxes. **(E)** shRNA knockdown of endogenous HNRNPC in the presence of V5-tagged WT or RRM-swapped exogenous HNRNPC, measured via western blot. Representative tubulin normalized total HNRNPC quantified by densitometry calculations shown at bottom. **(F)** Autoradiogram of V5-WT (top) or RRM swapped (bottom) HNRNPC, in the presence or absence of endogenous HNRNPC knockdown. Excised regions for sequencing indicated with red boxes.

### Figure S3

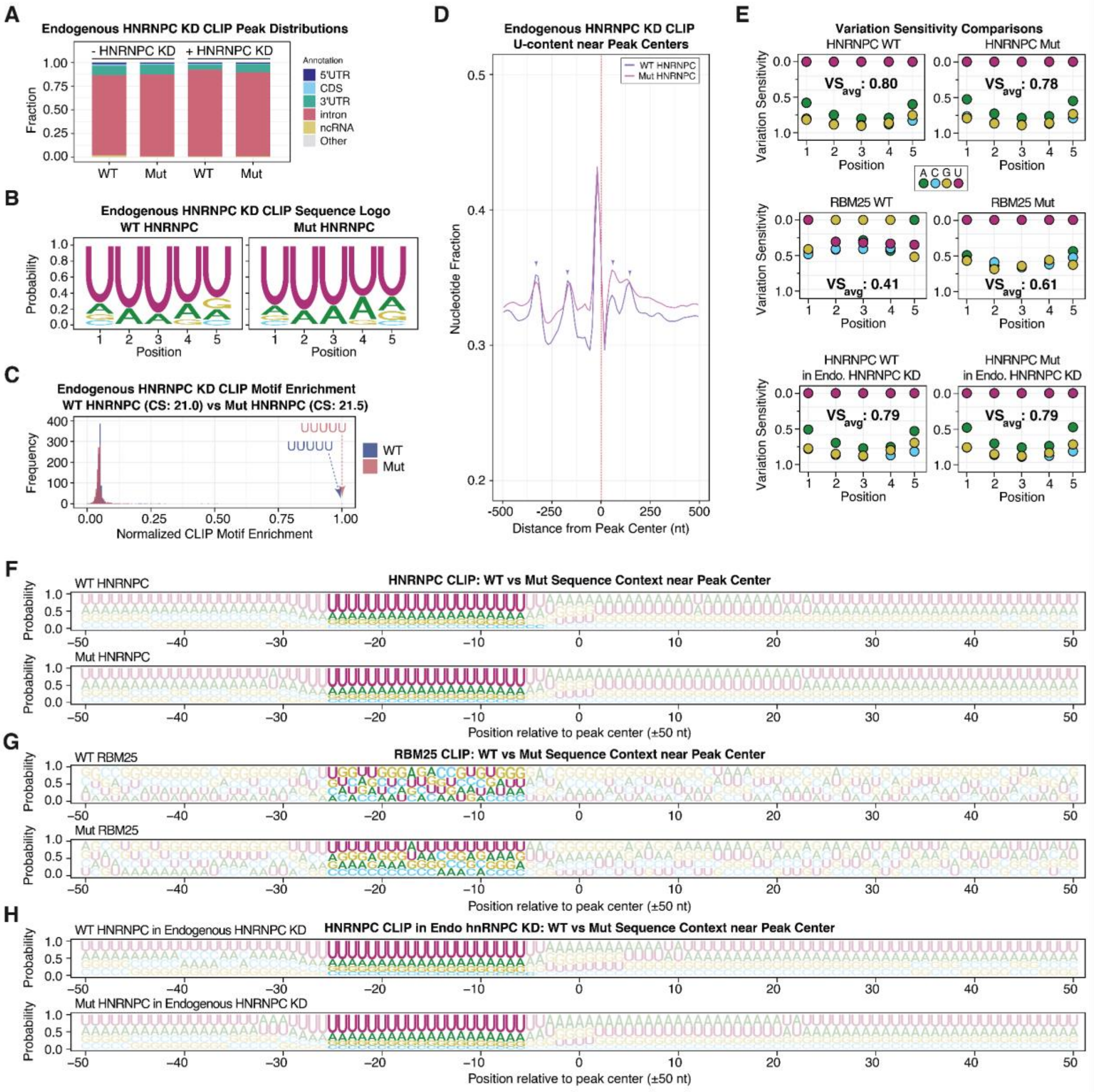


**Figure S3: Characterization of WT and mutant HNRNPC CLIP with endogenous HNRNPC knockdown. (A)** Genomic distribution of HNRNPC CLIP peaks in the presence (+) or absence (-) of endogenous HNRNPC knockdown (KD) for WT and mutant (Mut) proteins. **(B)** Sequence logos derived from motif enrichment for WT and Mut HNRNPC under HNRNPC KD conditions. **(C)** 5-mer motif enrichment distributions based on CLIP motif enrichment for WT and Mut HNRNPC with endogenous KD. Cellular specificity (CS) values are indicated. **(D)** U-content near CLIP peak centers for WT and Mut HNRNPC during endogenous KD. Periodic U-enrichment is indicated by purple markers. **(E)** Cellular variation sensitivity (CVS) plots and average CVS values ($VS_{avg}$) for all analyzed CLIP datasets, including HNRNPC and RBM25 WT/Mut pairs across different KD backgrounds. **(F)** Venn diagrams illustrating peak overlaps between: HNRNPC (in WT RBM25 OE) and WT RBM25 (left); WT RBM25 and Mut RBM25 (middle); and HNRNPC (in Mut RBM25 OE) and Mut RBM25 (right). **(G-J)** Sequence context logos showing nucleotide frequencies near peak centers (±50 nt) for the indicated CLIP datasets.

### Figure S4

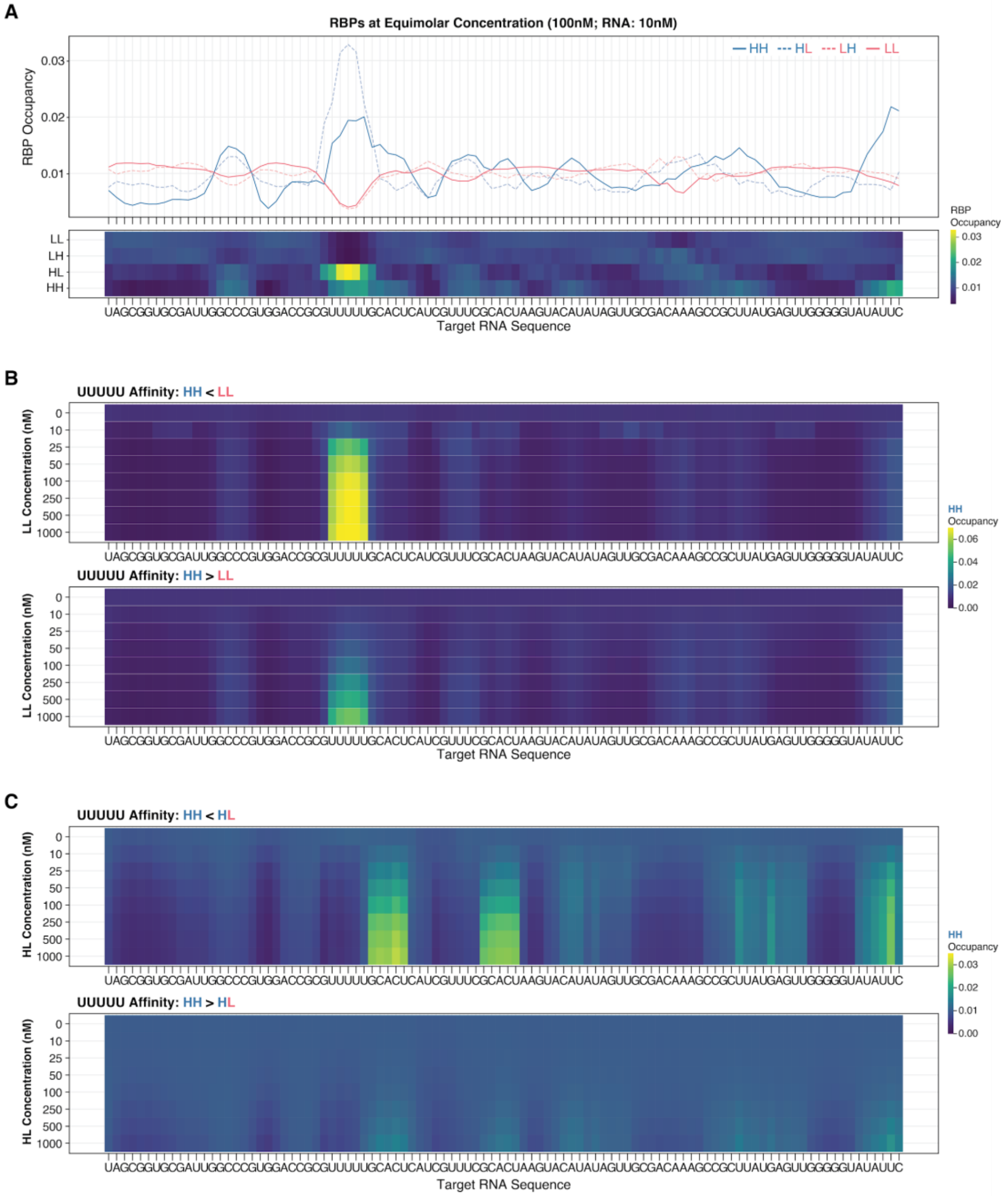


**Figure S4:** ***In silico*** **simulation results of RBP competitive binding on target RNA. (A)** Occupancies of the four model RBPs (HH, HL, LH, and LL) across the target RNA sequence. Simulations were performed with RBPs at 100 nM and RNA at 10 nM. The heatmap (bottom) summarizes the per-position occupancy values shown in the line plot (top). **(B)** HH occupancy across the target RNA in the presence of an LL competitor. Heatmaps illustrate competition across increasing competitor concentrations (0 to 1000 nM) for two scenarios: where the UUUUU affinity of HH is lower than LL (top) or higher than LL (bottom). **(C)** Same as **(B)** but depicting the competition between HH and an HL competitor.

Figure S5

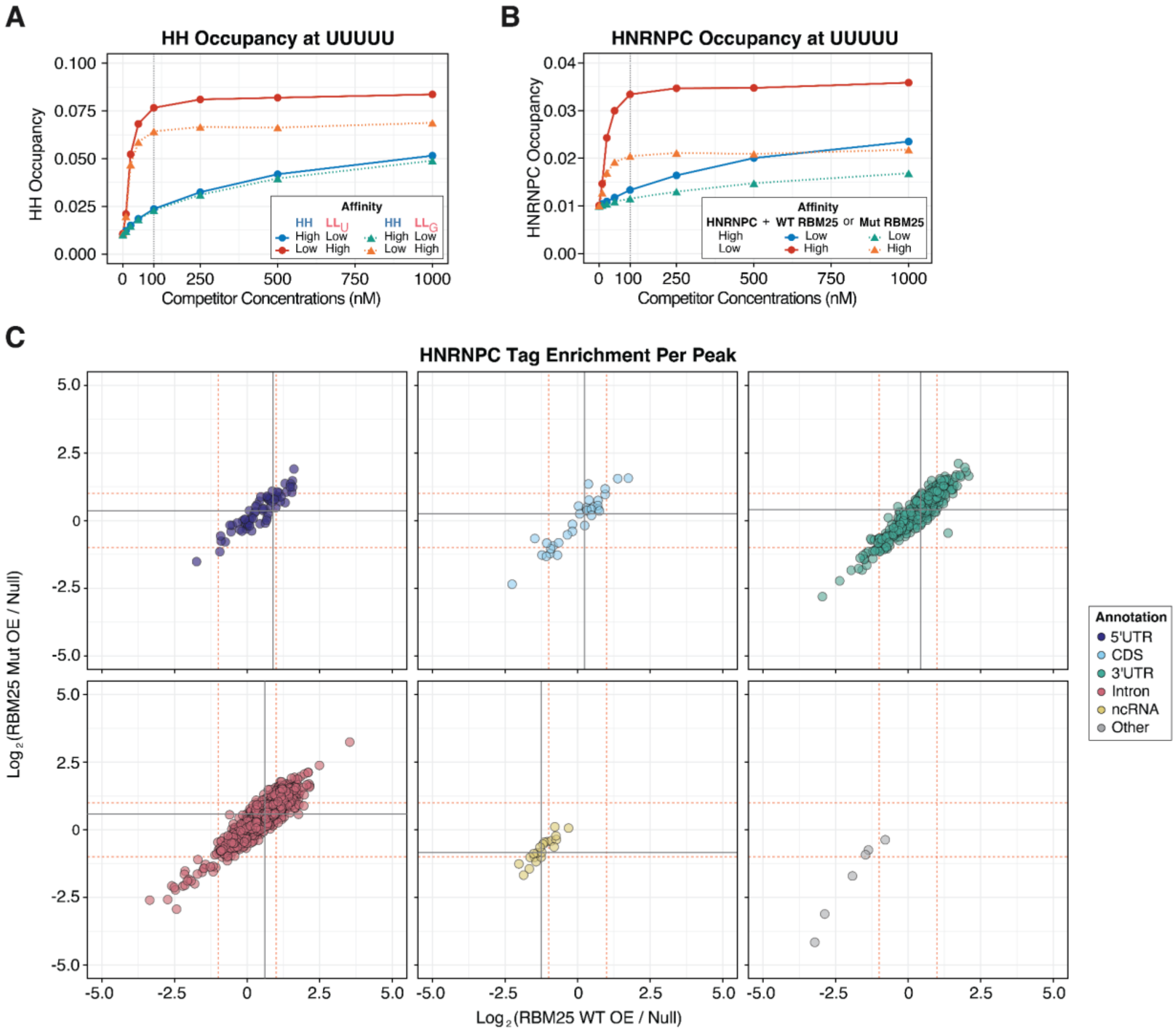


**Figure S5:** ***In silico*** **simulation and peak analysis results of Competition CLIP.** (**A**) *In silico* competition simulation between a high-specificity (HH) RBP and low-specificity competitors targeting either "UUUUU" or "GGGGG". (**B**) HNRNPC occupancy simulation at a target region using experimental CLIP motif enrichment data. **(C)** Per-peak HNRNPC tag enrichment comparison between WT and Mut RBM25 overexpression conditions by genomic annotation. Grey solid lines denote median enrichment values. Red dotted lines denote 2-fold enrichment.

**Figure S6**

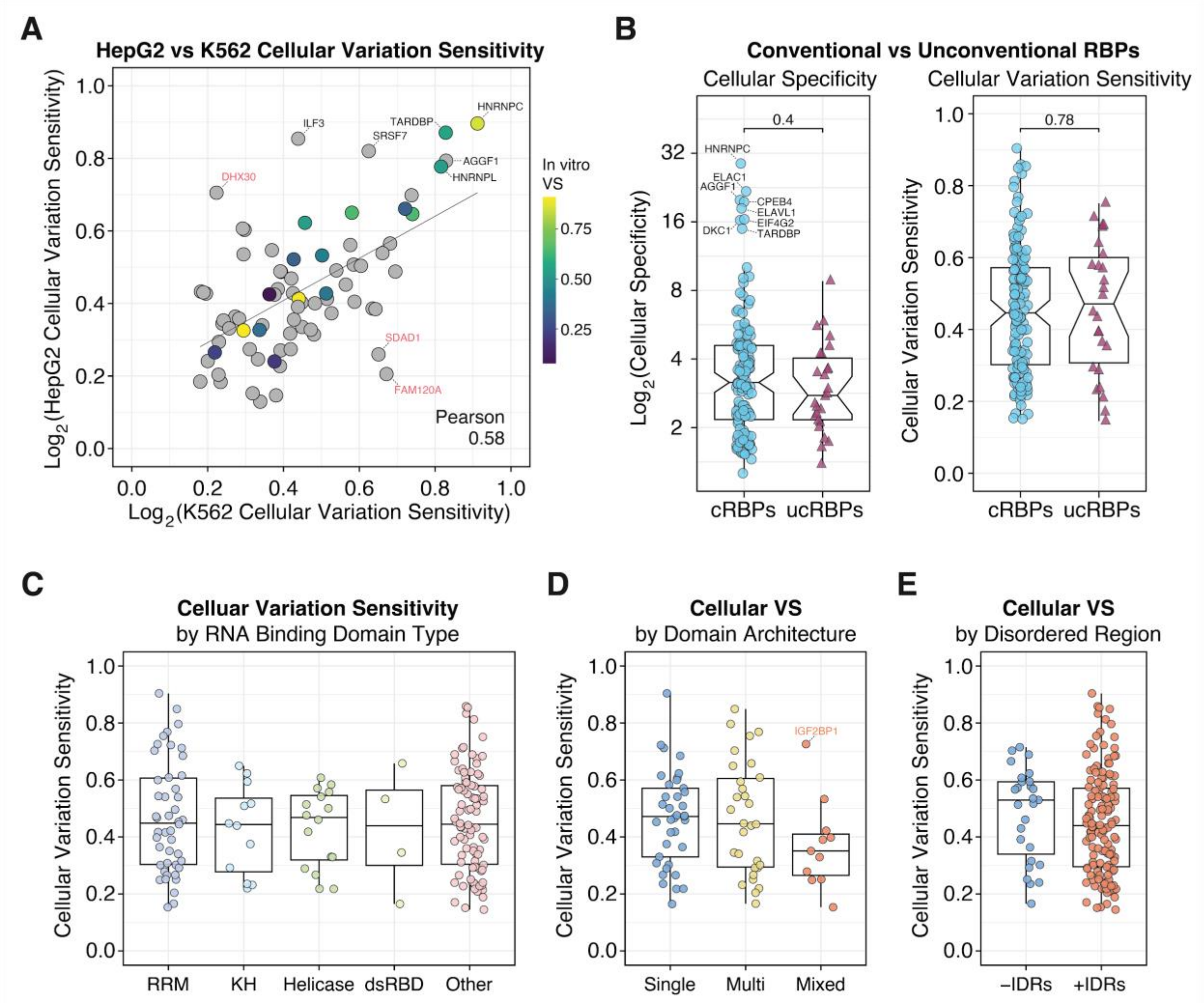


**Figure S6: Cellular variation sensitivity analysis across an RBP repertoire. (A)** Correlation between cellular variation sensitivity (CVS) values calculated from eCLIP data in HepG2 and K562 cell lines. Pearson r correlation is shown. RBPs in red are statistical outliers; those in black deviate from the linear trendline. **(B)** Cellular specificity (CS) and CVS distributions of conventional and unconventional RBPs. Outlier RBPs are annotated. **(C)** Distribution of CVS by RNA-binding domain (RBD) types. Selected outliers are annotated. **(D)** Distribution of CVS by domain architecture. **(E)** Distribution of CS by the presence of intrinsically disordered regions (IDRs).
